# Unifying the known and unknown microbial coding sequence space

**DOI:** 10.1101/2020.06.30.180448

**Authors:** Chiara Vanni, Matthew S. Schechter, Silvia G. Acinas, Albert Barberán, Pier Luigi Buttigieg, Emilio O. Casamayor, Tom O. Delmont, Carlos M. Duarte, A. Murat Eren, Robert D. Finn, Renzo Kottmann, Alex Mitchell, Pablo Sanchez, Kimmo Siren, Martin Steinegger, Frank Oliver Glöckner, Antonio Fernandez-Guerra

## Abstract

Genes of unknown function are among the biggest challenges in molecular biology, especially in microbial systems, where 40%-60% of the predicted genes are unknown. Despite previous attempts, systematic approaches to include the unknown fraction into analytical workflows are still lacking. Here, we propose a conceptual framework and a computational workflow that bridge the known-unknown gap in genomes and metagenomes. We showcase our approach by exploring 415,971,742 genes predicted from 1,749 metagenomes and 28,941 bacterial and archaeal genomes. We quantify the extent of the unknown fraction, its diversity, and its relevance across multiple biomes. Furthermore, we provide a collection of 283,874 lineage-specific genes of unknown function for *Cand*. Patescibacteria, being a significant resource to expand our understanding of their unusual biology. Finally, by identifying a target gene of unknown function for antibiotic resistance, we demonstrate how we can enable the generation of hypotheses that can be used to augment experimental data.

## Introduction

Thousands of isolate, single-cell, and metagenome-assembled genomes are guiding us towards a better understanding of life on Earth (Almeida et al., 2019; Cross et al., 2019; Delmont et al., 2020; Hug et al., 2016; Kopf et al., 2015; Pachiadaki et al., 2019; Pasolli et al., 2019; Sunagawa et al., 2015). At the same time, the ever-increasing number of genomes and metagenomes, unlocking uncharted regions of life’s diversity, (Brown et al., 2015; Eloe-Fadrosh et al., 2016; Hug et al., 2016) are providing new perspectives on the evolution of life (Parks et al., 2018; Spang et al., 2015). However, our rapidly growing inventories of new genes have a glaring issue: between 40% and 60% cannot be assigned to a known function (Almeida et al., 2020; Bernard et al., 2018; Carradec et al., 2018; Price et al., 2018). Current analytical approaches for genomic and metagenomic data (Chen et al., 2019; Franzosa et al., 2018; Huerta-Cepas et al., 2017; Mitchell et al., 2020; Quince et al., 2017) generally do not include this uncharacterized fraction in downstream analyses, constraining their results to conserved pathways and housekeeping functions (Quince et al., 2017). This inability to handle the unknown is an immense impediment to realizing the potential for discovery of microbiology and molecular biology at large (Bernard et al., 2018; Hanson et al., 2010). Predicting function from traditional single sequence similarity appears to have yielded all it can (Arnold, 2018, 1998; Brandenberg et al., 2017), thus several groups have attempted to resolve gene function by other means. Such efforts include combining biochemistry and crystallography (Jaroszewski et al., 2009); using environmental co-occurrence (Buttigieg et al., 2013); by grouping those genes into evolutionarily related families (Bateman et al., 2010; Brum et al., 2016; Wyman et al., 2018; Yooseph et al., 2007); using remote homologies (Bitard-Feildel and Callebaut, 2017; Lobb et al., 2015); or more recently using deep learning approaches (Bileschi et al., 2019; Liu, 2017). In 2018, Price et al. (Price et al., 2018) developed a high-throughput experimental pipeline that provides mutant phenotypes for thousands of bacterial genes of unknown function being one of the most promising methods to tackle the unknown. Despite their promise, experimental methods are labor-intensive and require novel computational methods that could bridge the existing gap between the genes with known and unknown function.

Here we present a conceptual framework and a computational workflow that closes the gap by connecting genomic and metagenomic data by the exploitation of groups of homologous genes, and facilitates the integration of genes of unknown function into ecological, evolutionary, and biotechnological investigations. The conceptual framework is based on the partitioning of the known and unknown fractions into four different categories that reflects the level of darkness (Figure 1A). The category “Known” (K) contains these sequences predicted to harbor domains of known function described by Pfam (Finn et al., 2016). Some sequences without Pfam domains of unknown function, like the ones encoding for small and intrinsically disordered proteins, can have homology to characterized proteins, hence we classify them as “Known without Pfam” (KWP). Finally, sequences that cannot be classified by either of these criteria, are classified as “Genomic Unknown” (GU) when they may be found in the genomes in reference databases; and “Environmental Unknown” (EU) when only have been observed in environmental samples. By contextualizing the different categories with information from several sources (Figure 1C), we provide an invaluable resource for the investigation of the genes with unknown function and boost the current methods for its experimental characterization.

**Figure 1:**
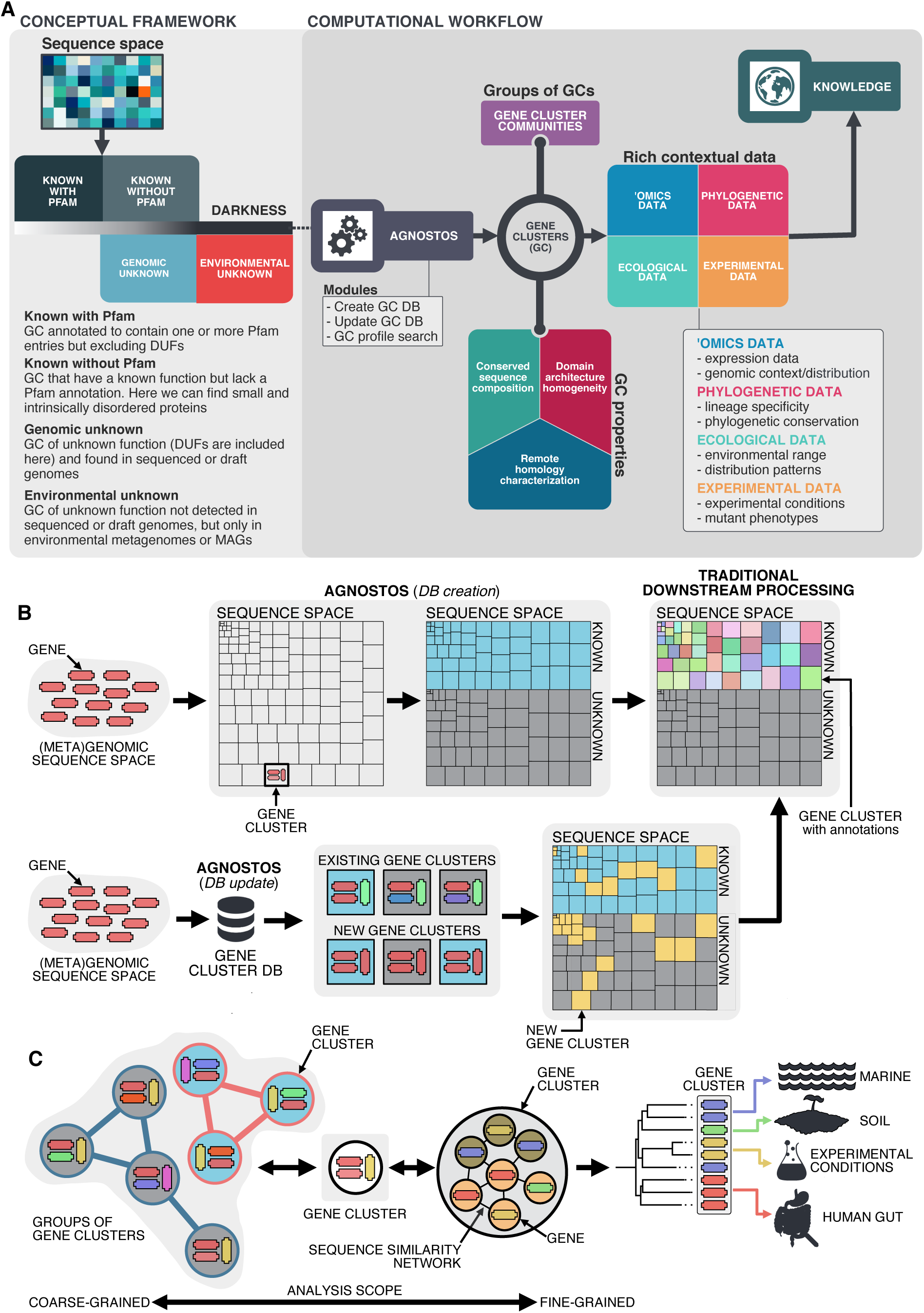
Conceptual framework to unify the known and unknown sequence space and integration of the framework in the current analytical workflows (A) Link between the conceptual framework and the computational workflow to partition the sequence space in the four conceptual categories. AGNOSTOS infers, validates and refines the GCs and combines them in gene cluster communities (GCCs). Then, it classifies them in one of the four conceptual categories based on their level of ‘darkness’. Finally, we add context to each GC based on several sources of information, providing a robust framework for generating hypotheses that can be used to augment experimental data. (B) The computational workflow provides two mechanisms to structure sequence space using GCs, de novo creation of the GCs (*DB creation*), or integrating the dataset in an existing GC database (*DB update*). The structured sequence space can then be plugged into traditional analytical workflows to annotate the genes within each GC of the known fraction. With AGNOSTOS, we provide the opportunity to integrate the unknown fraction into microbiome analyses easily. C) The versatility of the GCs enables analyses at different scales depending on the scope of our experiments. We can group GCs in gene cluster communities based on their shared homologies to perform coarse-grained analyses. On the other hand, we can design fine-grained analyses using the relationships between the genes in a GC, i.e., detecting network modules in the GC inner sequence similarity network. Additionally, given that GCs are conserved across environments, organisms and experimental conditions give us access to an unprecedented amount of information to design and interpret experimental data.

The application of our approach to 415,971,742 genes predicted from 1,749 metagenomes and 28,941 bacterial and archaeal genomes revealed that the unknown fraction (1) is smaller than has been previously reported, (2) is exceptionally diverse, and (3) is phylogenetically more conserved than the known fraction and predominantly lineage-specific at the Species level.

Finally, we show how we can connect all the outputs produced by our approach to augment the results from experimental data and add context to genes of unknown function through hypothesis-driven molecular investigations.

## Results

### A computational workflow to unify the known and the unknown coding sequence space

Driven by the concepts defined in the conceptual framework, we developed AGNOSTOS, a computational workflow that infers, validates, refines, and classifies groups of homologous genes or gene clusters (GCs) in the four proposed categories (Fig. 1A; Fig. 1B; Supp. Fig 1). AGNOSTOS produces GCs with a highly conserved intra-homogeneous structure (Fig. 1B), both in terms of sequence similarity and domain architecture homogeneity; it exhausts any existing homology to known genes and provides a proper delimitation of the unknown genes before classifying each GC in one of the four categories (Methods). In the last step, we decorate each GC with a rich collection of contextual data compiled from different sources or generated by analyzing the GC contents in different contexts (Fig. 1A). For each GC, we also offer several products that can be used for analytical purposes like improved representative sequences, consensus sequences, sequence profiles for MMseqs2 (Steinegger and Soding, 2017) and HHblits (Steinegger et al., 2019a), or the GC members as a sequence similarity network (Methods). To complement the collection, we also provide a subset of what we define as *high-quality* GCs. The defining criteria are (1) the representative is a complete gene and (2) more than one-third of genes within a GC are complete genes.

### Partitioning and contextualizing the coding sequence space of genomes and metagenomes

First, we used AGNOSTOS in a metagenomic context to show its ability to process complex and noisy datasets. We explored the unknown sequence space of 1,749 human and marine metagenomes. In total, we predicted 322,248,552 genes from the environmental dataset and assigned Pfam annotations to 44% of them (Supp. Fig. 2A). Next, AGNOSTOS clustered the predicted genes in 32,465,074 GCs and flagged those gene clusters that contain less than ten genes (Skewes-Cox et al., 2014). Flagged gene clusters are not processed through the validation workflow because they don’t have enough sequences to provide reliable results. We flagged 29,461,177 gene clusters, 19,911,324 of them being singletons (Supp. Fig. 2A; Supp. Note 1). A total of 3,003,897 GCs (83% of the original genes) will go through the validation steps. The validation process selected 2,940,257 *good-quality* clusters (Fig. 1B; Supp. Table 1; Supp. Note 2), which resulted in 43% of them being members of the unknowns after the classification and remote homology refinement steps (Supp. Fig. 2A, Supp. Note 3).

Lastly, we demonstrate how AGNOSTOS can integrate a new dataset into the already existing metagenomic database by integrating 28,941 genomes from the GTDB_r86 (Supp. Fig. 2A). With this integration we can build links between the environmental and genomic sequence space by expanding the final collection of GCs with the genes predicted from GTDB_r86. Surprisingly, the integration showed that the environmental GCs (human and marine) not only included a large proportion of the genes predicted from GTDB_r86 (72%) but also that most of these genes (90%) are part of the known sequence space (Supp. Fig. 2A; Supp. Note 4; Supp. Note 5). Only 22% of the GTDB_r86 genes created new GCs (2,400,037), being most of these genes (84%) classified as unknowns by AGNOSTOS (Supp. Fig. 2A; Supp. Note 4; Supp. Note 5). Finally, only a small proportion of the genes from GTDB_r86 (6%) resulted in singletons (Supp. Fig. 2A; Supp. Note 4; Supp. Note 5).

The final dataset after being validated and categorized resulted in 5,287,759 GCs (Supp. Fig. 2A), with both datasets sharing only 922,599 GCs (Supp. Fig. 2B). The integration of GTDB_r86 into the metagenomic database increased the proportion of GCs in the unknown sequence space from 43% to the 54%.

Additionally, AGNOSTOS identified a subset of 203,217 *high-quality* GCs (Supp. Table 2). In these *high-quality* GCs, we identified 12,313 clusters potentially encoding for small proteins (<= 50 amino acids). Most of these GCs are unknown (66%), which agrees with recent findings on novel small proteins from metagenomes (Sberro et al., 2019). We also observed that the KWP category contains the largest proportion of incomplete genes (Supp. Table 3), disrupting the detection and assignment of Pfam domains. But it also incorporates sequences with an unusual amino acid composition that have homology to proteins with high levels of disorder in the DPD database (Perdigão et al., 2017) and has characteristic functions of intrinsically disordered proteins (Habchi et al., 2014) (IDP) like cellular processes and signaling as predicted by eggNOG annotations (Supp. Table 4).

As part of the workflow, each GC is complemented with a rich set of information, as shown in Fig 1A (Supp. Table 5; Supp. Note 6).

### Beyond the twilight zone, communities of gene clusters

To find relationships between gene clusters, we grouped them in gene cluster communities (GCCs) (Fig. 2A) using remote homologies. As we are dealing with the unknown, we identified GCCs independently in each category, using the gene clusters from category K as a reference to identify the best parameters for the clustering of the gene cluster homology network (Methods; Fig. 2B; Supp. Note 7) and applying the learnt parameters to the other categories. The grouping reduced the final collection of GCs by 87%, producing 673,601 GCCs (Methods; Fig. 2B; Supp. Note 7).

**Figure 2:**
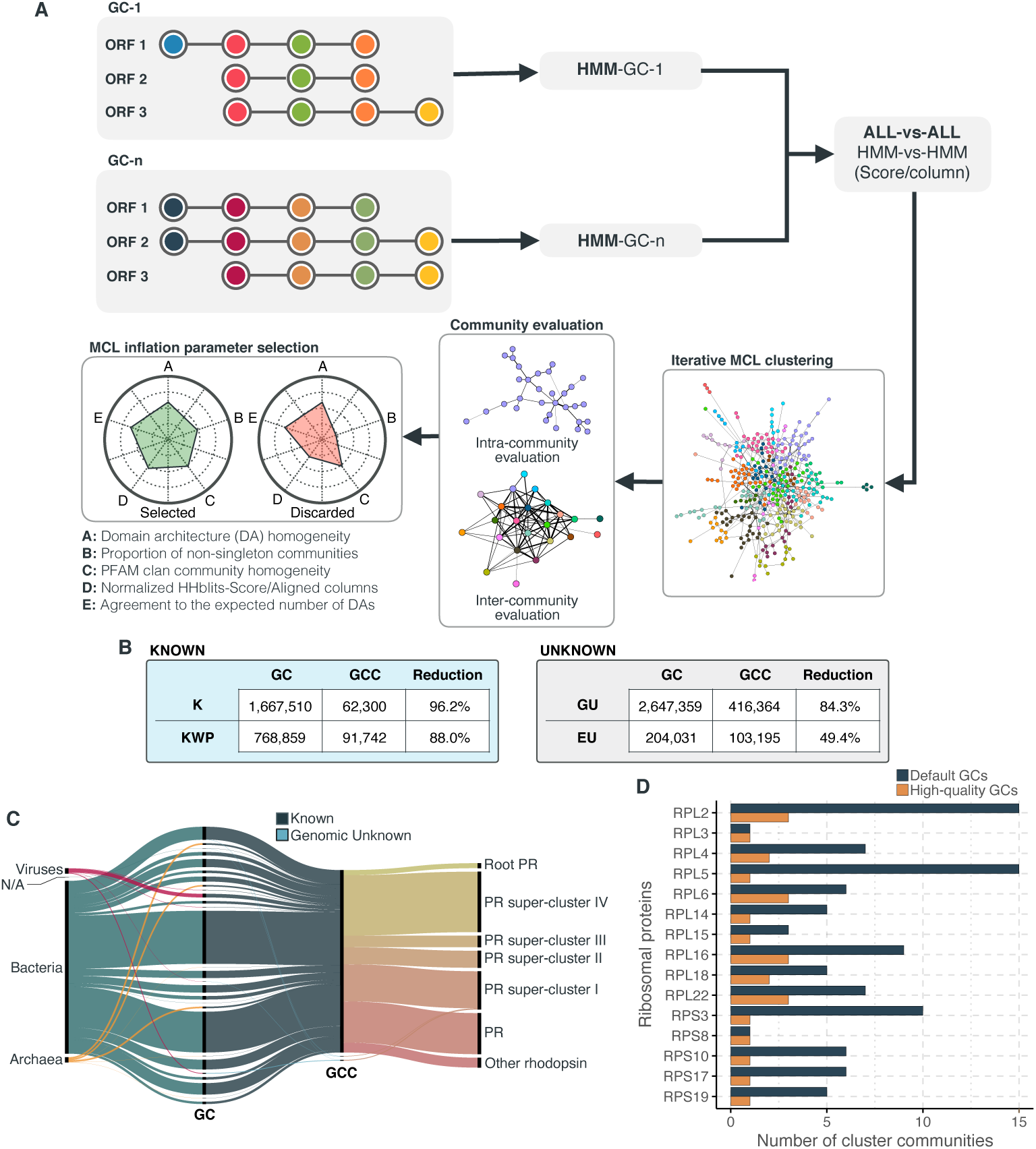
Overview and validation of the workflow to aggregate GCs in communities. (A) We inferred a gene cluster homology network using the results of an all-vs-all HMM gene cluster comparison with HHBLITS. The edges of the network are based on the HHblits-score/Aligned-columns. Communities are identified by an iterative screening of different MCL inflation parameters and evaluated using five different metrics that consider the inter- and intra-community properties. (B) Comparison of the number of GCs and GCCs for each of the functional categories. (C) Validation of the GCCs inference based on the environmental genes annotated as proteorhodopsins. Ribbons in the alluvial plot are genes, and each stacked bar corresponds (from left to right) to the (1) gene taxonomic classification at the domain level, (2) GC membership, (3) GCC membership and (4) MicRhoDE operational classification. (D) Validation of the GCCs inference based on ribosomal proteins based on standard and high-quality GCs.

We validated our approach to capture remote homologies between related GCs using two well-known gene families present in our environmental datasets, proteorhodopsins (Olson et al., 2018) and bacterial ribosomal proteins (Méheust et al., 2019). Theoretically, we would expect to have one or a very low number of GCCs for each of these gene families.

We identified 64 GCs (12,184 genes) and 3 GCCs (Supp. Note 8) containing genes classified as proteorhodopsin (PR). One community from the category K contained 99% of the PR annotated genes (Fig. 2C), except 85 genes taxonomically annotated as viral and assigned to the *PR Supercluster I* (Boeuf et al., 2015) within two GU communities (five GU gene clusters; Supp. Note 8).

For the ribosomal proteins, the results were not so satisfactory. We identified 1,843 GCs (781,579 genes) and 98 GCCs. The number of GCCs is larger than the expected number of ribosomal protein families (16) used for validation. When we used *high-quality* GCs (Supp. Note 8), we got closer to the expected number of GCCs (Fig. 2D). With this subset, we identified 26 GCCs and 145 GCs (1,687 genes). The cross-validation of our method against the approach used in Méheust et al. (Méheust et al., 2019) (Supp. Note 9) confirms the intrinsic complexity of analyzing metagenomic data. Both approaches showed a high agreement in the GCCs identified (Supp. Table 9-1). Still, our method inferred fewer GCCs for each of the ribosomal protein families (Supp. Fig. 9-3), coping better with the complexities of a metagenomic setup, such as incomplete genes (Supp. Table 6).

### A smaller but highly diverse unknown coding sequence space

By combining clustering and remote homology searches we reduced the extent of the unknown sequence space compared to what is reported by the traditional genomic and metagenomic analysis approaches where the unknown fraction can add up to 60% in marine metagenomes (Salazar et al., 2019) or up to 40% in human metagenomes (Thomas and Segata, 2019) (Fig. 3A). Our workflow recruited as much as 71% of genes in human-related metagenomic samples and 65% of the genes in marine metagenomes into the known sequence space. In both human and marine microbiomes, the genomic unknown fraction showed a similar proportion of genes (21%, Fig. 3A). The number of genes corresponding to EU gene clusters is higher in marine metagenomes; 12% of the genes are part of this GC category. We obtained a comparable result when we evaluated the genes from the GTDB_r86, 75% of bacterial and 64% of archaeal genes were part of the known sequence space. Archaeal genomes contained more unknowns than those from Bacteria, where 30% of the genes are classified as genomic unknowns in Archaea, and only 20% in Bacteria (Fig. 3A; Supp. Table 7).

**Figure 3:**
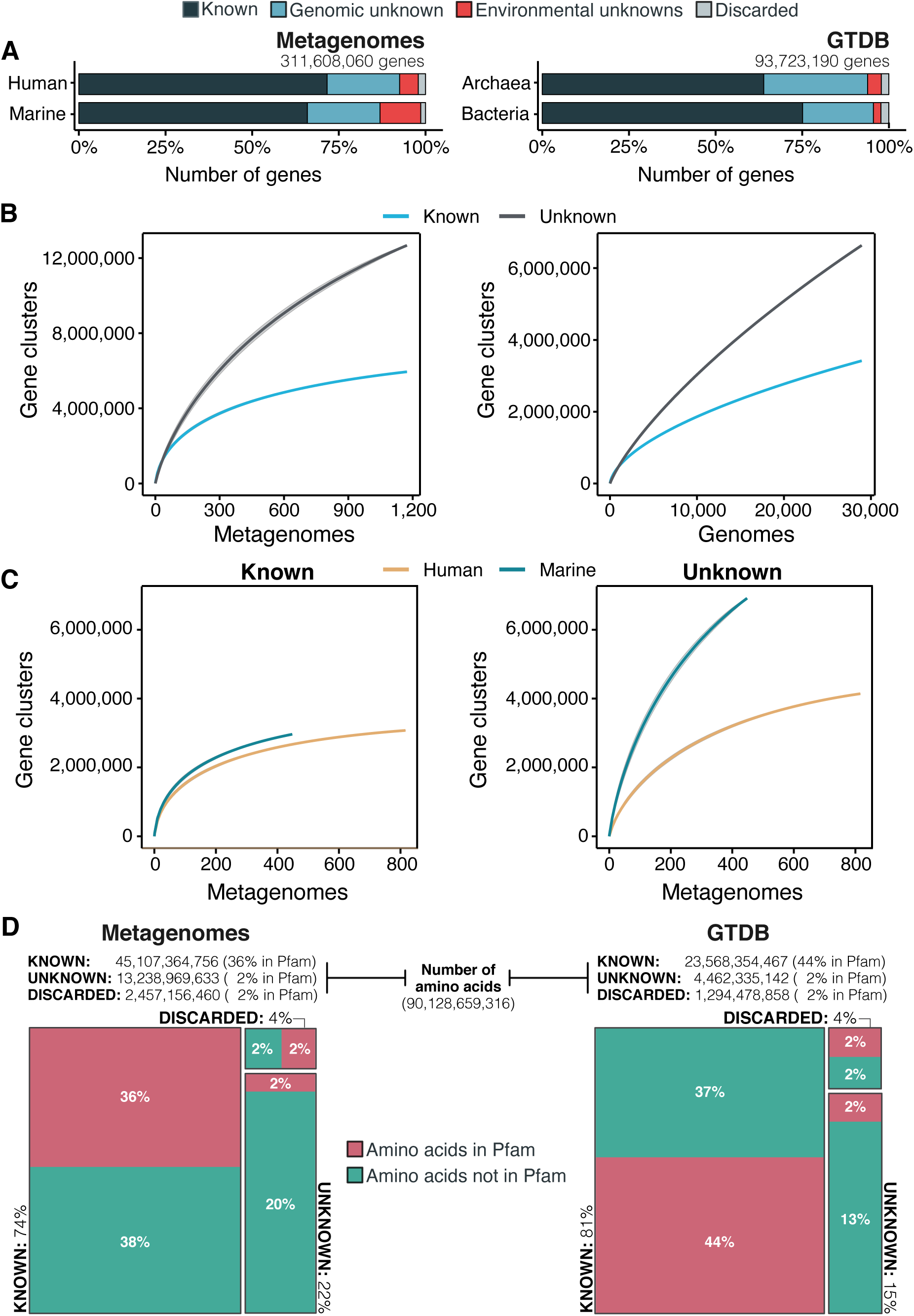
The extent of the known and unknown sequence space (A) Proportion of genes in the known and unknown. (B) Accumulation curves for the known and unknown sequence space at the GC-level for the metagenomic and genomic data. from TARA, MALASPINA, OSD2014 and HMP-I/II projects. (C) Collector curves comparing the human and marine biomes. Colored lines represented the mean of 1000 permutations and shaded in grey the standard deviation. Non-abundant singleton clusters were excluded from the accumulation curves calculation. (D) Amino acid distribution in the known and unknown sequence space.

To further evaluate the differences between the known and unknown sequence space, we calculated the accumulation rates of GCs and GCCs, using them as operational protein families (Schloss and Handelsman, 2008). For the metagenomic dataset we used 1,264 metagenomes (18,566,675 GCs and 282,580 GCCs) and for the genomic dataset 28,941 genomes (9,586,109 GCs and 496,930 GCCs). The rate of accumulation of unknown GCs was three times higher than the known (2X for the genomic), and in both cases the curves were far from reaching a plateau (Fig. 3B). This is not the case for the GCC accumulation curves (Supp. Fig. 4B), which reached a plateau.

The accumulation rate is largely determined by the number of singletons, especially singletons from EUs (Supp. note 10 and Supp. Fig. 5). While the accumulation rate of known GCs between marine and human metagenomes is almost identical, there are striking differences for the unknown GCs (Fig. 3C). These differences are maintained even when we remove the virus-enriched samples from the marine metagenomes (Supp. Fig. 4A). Although the marine metagenomes include a large variety of environments, from coastal to the deep sea, the known space remains quite constrained.

Next, we wanted to know how much of the sequence space we integrated with AGNOSTOS was found in other databases (Supp. Note 11). Despite only including marine and human metagenomes in our database, we already cover, in average, 76% of the sequence space of seven datasets spanning different databases and environments (Supp. Fig. 11-1). By screening MGnify (Mitchell et al., 2020) (release 2018_09; 11 biomes; 843,535,6116 proteins) we identified freshwater, soil and human non-digestive as the biomes less covered by our data (Supp. Fig. 6). Two of the seven analyzed datasets are designed to study genes of unknown function (Supp. Table 11-1). On (Wyman et al., 2018), where they defined Function Unknown Families of homologous proteins (FunkFams), we identified 20% of their FunkFams to be members of the known sequence space. On (Price et al., 2018), we classified as known, 44% of the genes of unknown function used in their experimental conditions.

One indirect consequence of our approach is that we can provide a detailed view of the sequence space at the amino acid level. We estimated the number of amino acids belonging to the known or unknown sequence space, and how many of these amino acids are contained within Pfam domain boundaries. From the 90,128,659,316 amino acids analyzed, most of the amino acids in metagenomes (74%) and genomes (80%) are in the known sequence space (Fig. 3D; Supp. Table 7) while only 22% in metagenomes and 15% in genomes are part of the unknowns. In both cases, approximately 40% of the amino acids in the known sequence space were part of a Pfam domain (Fig. 3D; Supp. Table 7). While this result is expected based on the large number of genes present in the known space (Fig. 3A), what is surprising is the low proportion of amino acids (2%) corresponding to DUF Pfam domains in genomes and metagenomes (Fig. 3D). If we use as reference the proportions of amino acids observed in the known sequence space, we can hypothesize that there are still many DUFs to be unearthed. With AGNOSTOS and its throughout validation and characterization of the genomic and environmental unknowns, we provide the basic building blocks (gene clusters’ multiple sequence alignments) to identify conserved regions that might become new potential DUFs.

### The unknown sequence space has a limited ecological distribution in human and marine environments

Although the role of the unknown fraction in the environment is still a mystery, the large number of gene counts and abundance observed underlines its inherent ecological relevance (Fig. 4A). In some metagenomes, the genomic unknown fraction can account for more than 40% of the total gene abundance observed (Fig. 4A). The environmental unknown fraction is also relevant in several samples, where singleton GCs are the majority (Fig. 4A). We identified two metagenomes with an unusual composition in terms of environmental unknown singletons. The marine metagenome corresponds to a sample from Lake Faro (OSD42), a meromictic saline with a unique extreme environment where Archaea plays an important role (La Cono et al., 2013). And the HMP metagenome (SRS143565) that corresponds to a human sample from the right cubital fossa from a healthy female subject. To understand this unusual composition, we should perform further analyses to discard potential technical artifacts like sample contamination.

**Figure 4:**
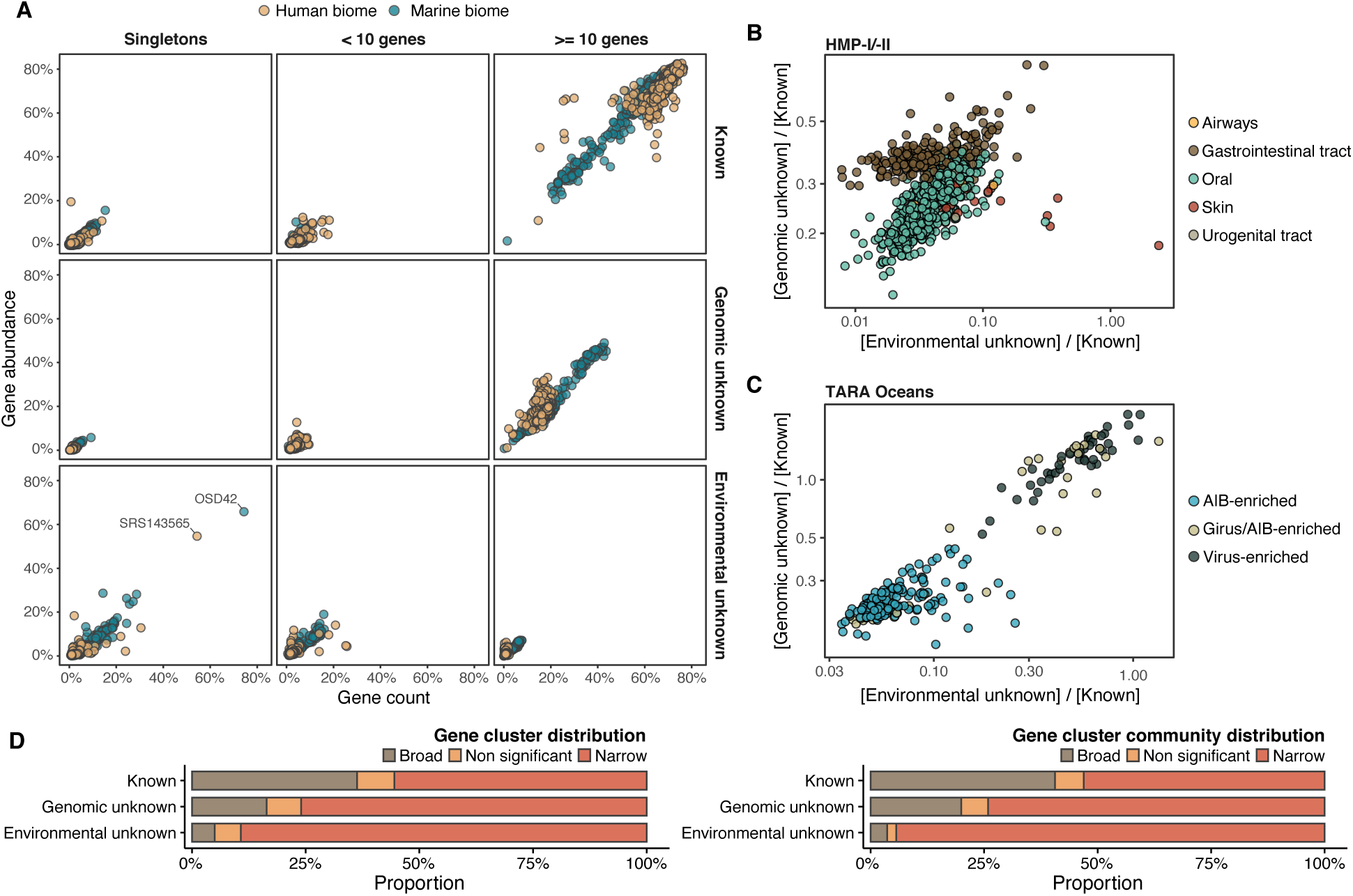
Distribution of the unknown sequence space in the human and marine metagenomes (A) Ratio between the proportion of the number of genes and their estimated abundances per cluster category and biome. Columns represented in the facet depicts three cluster categories based on the size of the clusters. (B) Relationship between the ratio of Genomic unknowns and Environmental unknowns in the HMP-I/II metagenomes. Gastrointestinal tract metagenomes are enriched in Genomic unknown sequences compared to the other body sites. (C) Relationship between the ratio of Genomic unknowns and Environmental unknowns in the TARA Oceans metagenomes. Girus and virus enriched metagenomes show a higher proportion of both unknown sequences (genomic and environmental) than the Archaea|Bacteria enriched fractions. (D) Environmental distribution of GCs and GCCs based on Levin’s niche breadth index. We obtained the significance values after generating 100 null gene cluster abundance matrices using the *quasiswap* algorithm.

The ratio between the unknown and known GCs is useful to reveal which metagenomes are enriched in GCs of unknown function (upper left quadrant in Fig. 4B-C) and it can be used as a proxy to assess the sequence contained in a metagenome. In human metagenomes, this ratio can distinguish between body sites, with the gastrointestinal tract, an ecologically complex environment (Qin et al., 2010), significantly enriched with genomic unknowns. Furthermore, it is not surprising that the human and marine metagenomes with the largest ratio of unknowns are those samples enriched with viral sequences. Specifically, in the HMP metagenomes are those samples identified to contain crAssphages (Dubinkina et al., 2016; Edwards et al., 2019) and HPV viruses (Ma et al., 2014) (Supp. Table 8; Supp. Fig. 7). While, in marine metagenomes (Fig. 4C) the highest ratio in genomic and environmental unknowns correspond to the ones enriched with viruses and giant viruses.

We performed a large-scale analysis to investigate the occurrence patterns of the GCs in the environment by analyzing their abundance and distribution breadth. The narrow distribution of the unknown fraction (Fig. 4D) suggests that these GCs might provide a selective advantage and be necessary to adapt to specific environmental conditions. But the pool of broadly distributed environmental unknowns is the most exciting result. We identified traces of potential ubiquitous organisms left uncharacterized by traditional approaches, as more than 80% of these GCs cannot be associated with a metagenome-assembled genome (MAG) (Supp. Table 9, Supp. Note 12).

### The genomic unknown coding sequence space is lineage-specific

With the inclusion of the genomes from GTDB_r86, we have access to a phylogenomic framework that can be used to assess how exclusive is a GC within a lineage (lineage-specifity (Mendler et al., 2019)) and how conserved (phylogenetic conservation) is a GC across the different clades in the GTDB_r86 phylogenomic tree (Martiny et al., 2013). We identified 781,814 lineage-specific GCs and 464,923 phylogenetically conserved (P < 0.05) GCs in Bacteria (Supp. Table 10; Supp. Note 13 for Archaea). The number of lineage-specific GCs increases with the Relative Evolutionary Distance (Parks et al., 2018) (Fig. 5A) and differences between the known and the unknown fraction start to be evident at the Family level resulting in 4X more unknown lineage-specific GCs at the Species level. In general terms, the unknown GCs are more phylogenetically conserved (GCs shared among members of deep clades) than the known (Fig. 5B, p < 0.0001), revealing the importance of the genome’s uncharacterized fraction. However, the lineage-specific unknown GCs are less phylogenetically conserved (Fig. 5B) than the known, agreeing with the large number of lineage-specific GCs observed at Genus and Species level (Fig. 5A).

**Figure 5:**
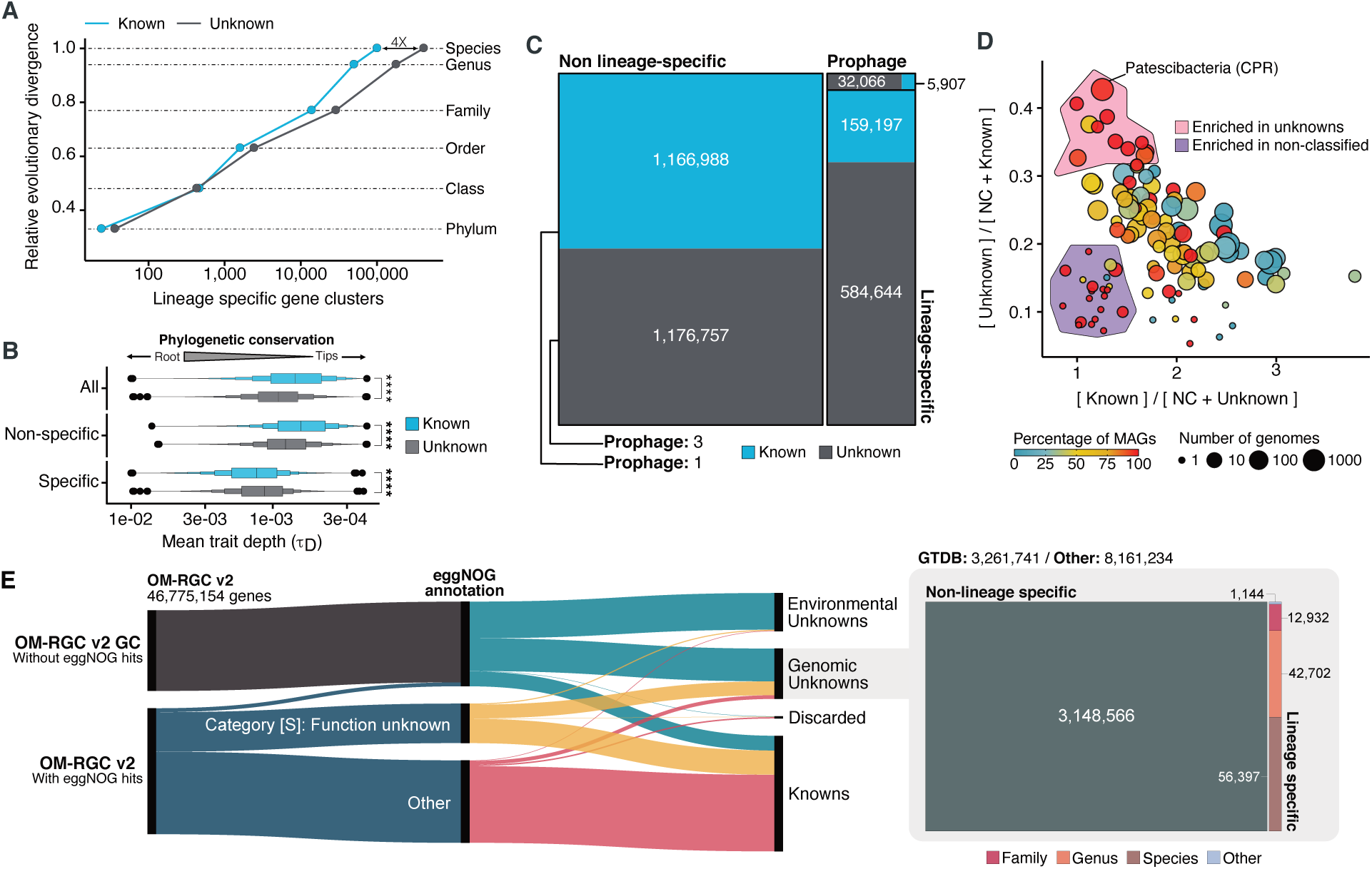
Phylogenomic exploration of the unknown sequence space. (A) Distribution of the lineage-specific GCs by taxonomic level. Lineage-specific unknown GCs are more abundant in the lower taxonomic levels (Genus, Species). (B) Phylogenetic conservation of the known and unknown sequence space in 27,372 bacterial genomes from GTDB_r86. We observe differences in the conservation between the known and the unknown sequence space for lineage- and non-lineage specific GCs (paired Wilcoxon rank-sum test; all p-values < 0.0001). (C) The majority of the lineage-specific clusters are part of the unknown sequence space, being a small proportion found in prophages present in the GTDB_r86 genomes. (D) Known and unknown sequence space of the 27,732 GTDB_r86 bacterial genomes grouped by bacterial phyla. Phyla are partitioned based on the ratio of known to unknown GCs and vice versa. Phyla enriched in MAGs have higher proportions in GCs of unknown function. Phyla with a high proportion of non-classified clusters (NC; discarded during the validation steps) tend to contain a small number of genomes. (E) The alluvial plot’s left side shows the uncharacterized (OM-RGC v2 GC) and characterized (OM-RGC v2) fraction of the gene catalog. The functional annotation is based on the eggNOG annotations provided by Salazar et al.(Salazar et al., 2019). The right side of the alluvial plot shows the new organization of the OM-RGC v2 sequence space based on the approach described in this study. The treemap in the right links the metagenomic and genomic space adding context to the unknown fraction of the OM-RGC v2

One potential confounding factor that might contribute to inflate the number of lineage-specific GCs in the unknown fraction, is the presence of prophages owing to their potential host specificity (Ross et al., 2016). To discard the possibility that the lineage-specific GCs of unknown function have a viral origin, we screened all GTDB_r86 genomes for prophages. We only found 37,163 lineage-specific GCs in prophage genomic regions, being 86% of these GCs of unknown function.

After unveiling the potential relevance of the GCs of unknown function in bacterial genomes, we identified phyla in GTDB_r86 enriched with these types of clusters. A clear pattern emerged when we partitioned the phyla based on the ratio of known to unknown GCs and vice versa (Fig. 5D), the phyla with a larger number of MAGs are enriched in GCs of unknown function (Fig. 5D). Phyla with a high proportion of non-classified GCs (those discarded during the validation steps) contain a small number of genomes and are primarily composed of MAGs. These groups of phyla highly enriched in unknowns and represented mainly by MAGs include newly described phyla such as *Cand*. Riflebacteria and *Cand*. Patescibacteria (Anantharaman et al., 2018; Brown et al., 2015; Rinke et al., 2013), both with the largest unknown to known ratio.

One of the strengths of AGNOSTOS is the possibility of bridging genomic and metagenomic data and simultaneously unifying the known and unknown sequence space by integrating the new Ocean Microbial Reference Gene Catalog (Salazar et al., 2019) (OM-RGC v2) into our database. We assigned 26,170,875 genes to known GCs, 11,422,975 to genomic unknowns, 8,661,221 to environmental unknown and 520,083 were discarded. From the 11,422,975 genes classified as genomic unknowns, we could associate 3,261,741 to a GTDB_r86 genome and we identified 113,175 as lineage-specific. The alluvial plot in Fig. 5E depicts the new organization of the OM-RGC v2 after being integrated into our framework and how we can provide context to the two original types of unknowns in the OM-RGC (those annotated as category S in eggNOG (Huerta-Cepas et al., 2019) and those without known homologs in the eggNOG database (Salazar et al., 2019)) that can lead to potential experimental targets at the organism level to complement the metatranscriptomic approach proposed by Salazar et al. (Salazar et al., 2019).

### A structured sequence space augments the interpretation of experimental data

We selected one of the experimental conditions tested in Price et al. (Price et al., 2018) to demonstrate the potential of our approach to augment experimental data. We compared the fitness values in plain rich medium with added Spectinomycin dihydrochloride pentahydrate to the fitness in plain rich medium (LB) in *Pseudomonas fluorescens FW300-N2C3* (Fig. 6A). This antibiotic inhibits protein synthesis and elongation by binding to the bacterial 30S ribosomal subunit and interferes with the peptidyl tRNA translocation. We identified the gene with locus id AO356_08590 that presents a strong phenotype (fitness = -3.1; t = -9.1) and has no known function. This gene belongs to the genomic unknown GC GU_19737823. We can track this GC into the environment and explore the occurrence in the different samples we have in our database. As expected, the GC is mostly found in non-human metagenomes (Fig. 6B) as *Pseudomonas* are common inhabitants of soil and water environments (Heffernan et al., 2009). However, finding this GC also in human-related samples is very interesting due to the potential association of *P. fluorescens* and human disease where Crohn’s disease patients develop serum antibodies to this microbe (Scales et al., 2014). We can add another layer of information to the selected GC by looking at the associated remote homologs in the GCC GU_c_21103 (Fig. 6C). We identified all the genes in the GTDB_r86 genomes that belong to the GCC GU_c_21103 (Supp. Table 11) and explored their genomic neighborhoods. All members from GU_c_21103 are constrained to the class *Gammaproteobacteria*, and interestingly GU_19737823 is mostly exclusive to the order *Pseudomonadales*. The gene order in the different genomes analyzed is highly conserved, finding GU_19737823 after the *rpsF*::*rpsR* operon and before *rpll. rpsF* and *rpsR* encode for 30S ribosomal proteins, the prime target of spectinomycin. The combination of the experimental evidence and the associated data inferred by our approach provides strong support to generate the hypothesis that the gene AO356_08590 might be involved in the resistance to Spectinomycin.

**Figure 6:**
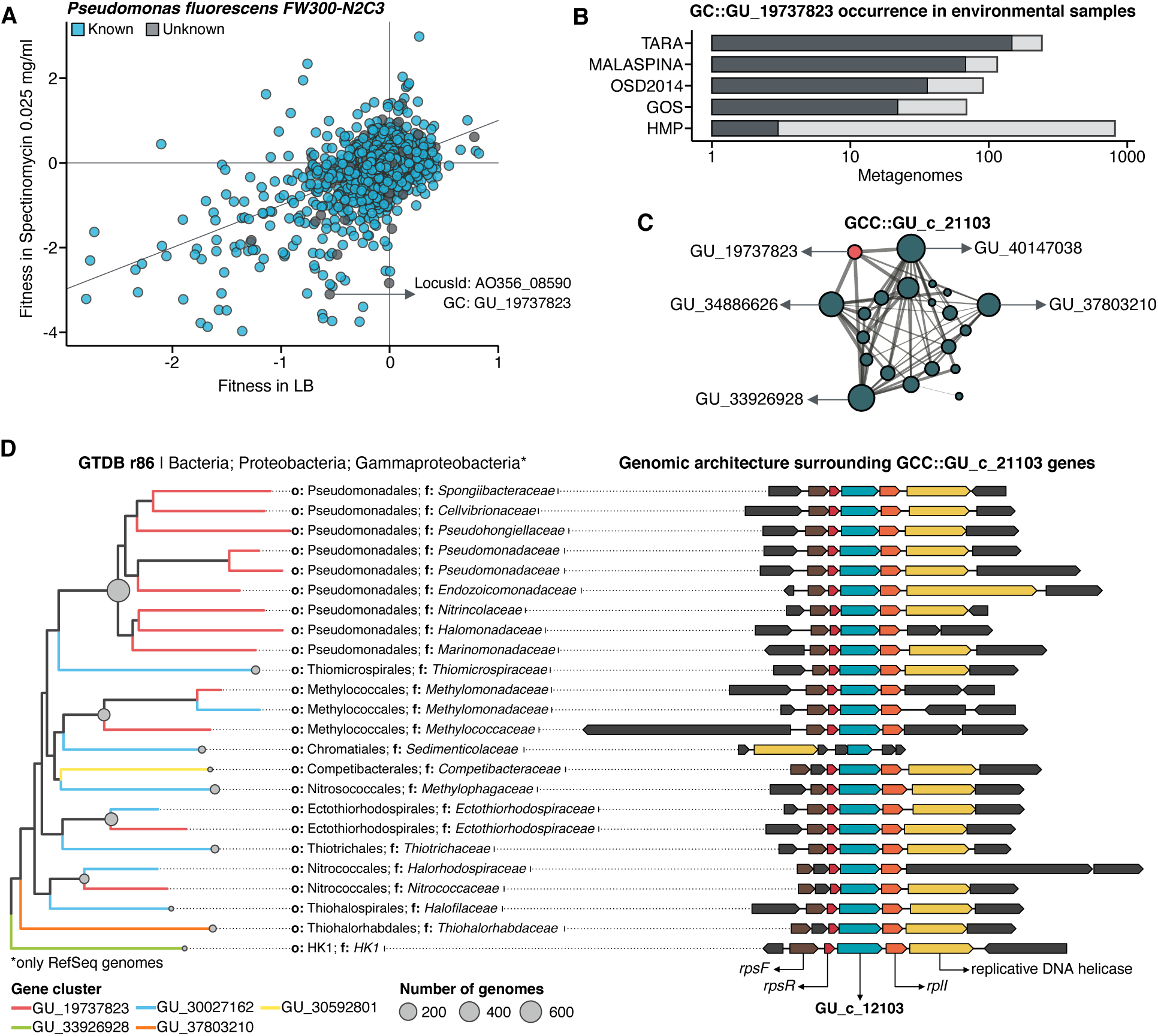
Augmenting experimental data with GCs of unknown function. (A) We used the fitness values from the experiments from Price et al. (Price et al., 2018) to identify genes of unknown function that are important for fitness under certain experimental conditions. The selected gene belongs to the genomic unknown GC GU_19737823 and presents a strong phenotype (fitness = -3.1; t = -9.1) (B) Occurrence of GU_19737823 in the metagenomes used in this study. Darker bars depict the number of metagenomes where the GC is found. (C) GU_19737823 is a member of the GCC GU_c_21103. The network shows the relationships between the different GCs members of the gene cluster community GU_c_21103. The size of the node corresponds to the node degree of each GC. Edge thickness corresponds to the bitscore/column metric. Highlighted in red is GU_19737823. (D) We identified all the genes in the GTDB_r86 genomes that belong to the GCC GU_c_21103 and explored their genomic neighborhoods. GU_c_21103 members were constrained to the class *Gammaproteobacteria*, and GU_19737823 is mostly exclusive to the order *Pseudomonadales*. The gene order in the different genomes analyzed is highly conserved, finding GU_19737823 after the *rpsF::rpsR* operon and before *rpll. rpsF* and *rpsR* encode for the *30S ribosomal protein S6* and *30S ribosomal protein S18*, respectively. The GTDB_r86 subtree only shows RefSeq genomes. Branch colors correspond to the different GCs found in GU_c_21103. The bubble plot depicts the number of genomes with a gene that belongs to GU_c_21103.

## Discussion

We present a new conceptual framework and computational workflow to unify the known and unknown sequence space. Using this framework, we performed an in-depth exploration of the microbial unknown sequence space, demonstrating that we can link the unknown fraction of metagenomic studies to specific genomes and provide a powerful tool for hypothesis generation. The framework introduces a subtle change of paradigm compared to traditional approaches where our objective is to provide the best representation of the unknown space. We gear all our efforts towards finding sequences without any evidence of known homologies by pushing the search space beyond the *twilight zone* of sequence similarity (Rost, 1999). With this objective in mind, we use gene clusters instead of genes as the fundamental unit to compartmentalize the sequence space owing to their unique properties (Fig. 1B). Gene clusters (1) provide a structured sequence space that helps to reduce its complexity, (2) are independent of the known and unknown fraction, (3) are conserved across environments and organisms, and can be used to aggregate information from different sources (Fig. 1A). Moreover, GCs provide a good compromise in terms of resolution for analytical purposes, and owing to their unique properties, one can perform analyses at different scales. For fine-grained analyses, we can exploit the gene associations within each GC; and for coarse-grained analyses, we can create groups of GCs based on their shared homologies (Fig. 1B).

AGNOSTOS integrates transparently into the standard operating procedure for analyzing metagenomes (Quince et al., 2017) adopted by the microbiome community. It can briefly be summarized into (1) assembly, (2) gene prediction, (3) gene catalog inference, (4) binning, and characterization. AGNOSTOS exploits recent computational developments (Steinegger and Söding, 2018; Steinegger and Soding, 2017) to maximize the information used when analyzing genomic and metagenomic data. In addition, we provide a mechanism to reconcile top-down and bottom-up approaches, thanks to the well-structured sequence space proposed by our framework. AGNOSTOS can create environmental- and organism-specific variations of a seed database based on gene clusters. Then, it integrates the predicted genes from new genomes and metagenomes and dynamically creates and classifies new GCs with those genes not integrated during the initial step (Fig. 1B). Afterward, the potential functions of the known GCs can be carefully characterized by incorporating them into the traditional standard operating procedure described previously.

One of the most appealing characteristics of our approach is that the GCs provide unified groups of homologous genes across environments and organisms indifferently if they belong to the known or unknown sequence space, and we can contextualize the unknown fraction using this genomic and environmental information. Our combination of partitioning and contextualization features a smaller unknown sequence space than previously reported (Steinegger et al., 2019b). On average, only 30% of the genes fall in the unknown fraction for our genomic and metagenomic data. One hypothesis to reconcile this surprising finding is that the methodologies to identify remotely homologous sequences in large datasets were computationally prohibitive until recently. New methods (Steinegger et al., 2019a; Steinegger and Soding, 2017), like the ones used in AGNOSTOS, enable large-scale remote homology searches. Still, one must apply conservative measures to control the trade-off between specificity and sensitivity to avoid overclassification.

We found that most of the sequence space at the gene and amino acid level is known, both in genomes and metagenomes. However, the GC accumulation curves showed that the unknown fraction is far more diverse than the known. When we combine the high diversity and its narrow ecological distribution, we can unveil the magnitude of the untapped unknown functional fraction and its potential importance for niche adaptation. In a genomic context and after ruling out the effect of prophages, the unknown fraction is predominantly Species’ lineage-specific and phylogenetically more conserved than the known fraction, supporting the signal observed in the environmental data emphasizing that we should not ignore the unknown fraction. It is worth noting that the high diversity observed in the unknowns only represents the 20% of the amino acids in the sequence space we analyzed, and only 10% of this unknown amino acid space is part of a Pfam domain (DUF and others). This contrasts with the numbers observed in the known sequence space, where Pfam domains include 50% of the amino acids. All this evidence combined strengthens the hypothesis that the genes of unknown function, especially the lineage-specific ones, might be associated with the mechanisms of microbial diversification and niche adaptation due to the constant diversification of gene families and the survival of new gene lineages (Francino, 2012; Muller, 2019).

Metagenome-assembled genomes are not only unveiling new regions of the microbial universe (42% of the genomes in GTDB_r86), but they are also enriching the tree of life with genes of unknown function. One excellent example is *Cand*. Patescibacteria, more commonly known as Candidate Phyla Radiation (CPR), a phylum that has raised considerable interest due to its unusual biology (Brown et al., 2015). We performed an in-depth exploration of this phylum, and we provide a collection of 54,343 lineage-specific GCs of unknown function at different taxonomic level resolutions (Supp. Table 12; Supp. Note 14), which will be a valuable resource for the CPR advancement research efforts.

One of the ultimate goals of our approach is to provide a mechanism to unlock the large pool of likely relevant data that remains untapped to analysis and discovery and boost insights from model organism experiments. We demonstrated the value of our approach by identifying a potential target gene of unknown function for antibiotic resistance. But severe challenges remain, such as the dependence on the quality of the assemblies and their gene predictions, as shown by the analysis of the ribosomal protein GCCs where many of the recovered genes are incomplete. While sequence assembly has been an active area of research (Roumpeka et al., 2017), this has not been the case for gene prediction methods (Roumpeka et al., 2017), which are becoming outdated (Ivanova et al., 2014) and cannot cope with the current amount of data. Alternatives like protein-level assembly (Steinegger et al., 2019b) combined with exploring the assembly graphs’ neighborhoods (Brown et al., 2020) become very attractive for our purposes. In any case, we still face the challenge of discriminating between genuine and artifactual singletons (Höps et al., 2018). There are currently no methods available to provide a plausible solution and, at the same time, being scalable. We devise a potential solution in the recent developments in unsupervised deep learning methods where they use large corpora of proteins to define a language model *embedding* for protein sequences (Heinzinger et al., 2019). These models could be applied to predict *embeddings* in singletons, which could be clustered or used to determine their coding potential. Another issue is that we might be creating more GCs than expected. We follow a conservative approach to avoid mixing multi-domain proteins in GCs owing to the fragmented nature of the metagenome assemblies that could result in the split of a GC. However, not only splitting can be a problem, but also lumping unrelated genes or GCs owing to the use of remote homologies. Although the inference of GCCs is using very sensitive methods to compare profile HMMs, low sequence diversity in GCs can limit its effectiveness.

Moreover, our approach is affected by the presence and propagation of contamination in reference databases, a significant problem in ‘omics (Breitwieser et al., 2019; Steinegger and Salzberg, 2020). In our case, we only use Pfam (Finn et al., 2016) as a source for annotation owing to its high-quality and manual curation process. The categorization process of our GCs depends on the information from other databases, and to minimize the potential impact of contamination, we apply methods that weight the annotations of the identified homologs to discriminate if a GC belongs to the known or unknown sequence space.

The results presented here prove that the integration and the analysis of the unknown fraction are possible. We are unveiling a brighter future, not only for microbiome analyses but also for boosting eukaryotic-related studies, thanks to the increasing number of projects, including metatranscriptomic data (Delmont et al., 2020; Vorobev et al., 2020). Furthermore, our work lays the foundations for further developments of clear guidelines and protocols to define the different levels of unknown (Thomas and Segata, 2019) and should encourage the scientific community for a collaborative effort to tackle this challenge.

## Material and methods

### Genomic and metagenomic dataset

We used a set of 583 marine metagenomes from four of the major metagenomic surveys of the ocean microbiome: Tara Oceans expedition (TARA) (Sunagawa et al., 2015), Malaspina expedition (Duarte, 2015), Ocean Sampling Day (OSD) (Kopf et al., 2015), and Global Ocean Sampling Expedition (GOS) (Rusch et al., 2007). We complemented this set with 1,246 metagenomes obtained from the Human Microbiome Project (HMP) phase I and II (Lloyd-Price et al., 2017). We used the assemblies provided by TARA, Malaspina, OSD and HMP projects and the long Sanger reads from GOS (Sanger et al., 1977). A total of 156M (156,422,969) contigs and 12.8M long-reads were collected (Supp. Table 6).

For the genomic dataset, we used the 28,941 prokaryotic genomes (27,372 bacterial and 1,569 archaeal) from the Genome Taxonomy Database (Parks et al., 2018) (GTDB) Release 03-RS86 (19th August 2018).

### Computational workflow development

We implemented a computation workflow based on Snakemake (Köster, 2018) for the easy processing of large datasets in a reproducible manner. The workflow provides three different strategies to analyze the data. The module *DB-creation* creates the gene cluster database, validates and partitions the gene clusters (GCs) in the main functional categories. The module *DB-update* allows the integration of new sequences (either at the contig or predicted gene level) in the existing gene cluster database. In addition, the workflow has a *profile-search* function to quickly screen samples using the gene cluster PSSM profiles in the database.

### Metagenomic and genomic gene prediction

We used Prodigal (v2.6.3) (Hyatt et al., 2010) in metagenomic mode to predict the genes from the metagenomic dataset. For the genomic dataset, we used the gene predictions provided by Annotree (Mendler et al., 2019), since they were obtained, consistently, with Prodigal v2.6.3. We identified potential spurious genes using the *AntiFam* database (Eberhardt et al., 2012). Furthermore, we screened for ‘*shadow*’ genes using the procedure described in Yooseph et al. (Yooseph et al., 2008).

### PFAM annotation

We annotated the predicted genes using the *hmmsearch* program from the *HMMER* package (version: 3.1b2) (Finn et al., 2011) in combination with the Pfam database v31 (Finn et al., 2016). We kept the matches exceeding the internal gathering threshold and presenting an independent e-value < 1e-5 and coverage > 0.4. In addition, we considered multi-domain annotations, and we removed overlapping annotations when the overlap is larger than 50%, keeping the ones with the smaller e-value.

### Determination of the gene clusters

We clustered the metagenomic predicted genes using the cascaded-clustering workflow of the MMseqs2 software (Steinegger and Söding, 2018) (“*--cov-mode 0 -c 0*.*8 --min-seq-id 0*.*3”*). We discarded from downstream analyses the singletons and clusters with a size below a threshold identified after applying a modification of the broken-stick model (Macarthur, 1957). We randomly split the number of gene clusters into p subsets, where p is defined by the proportion of outlier genes per gene cluster. The subsets are then sorted by decreasing size. We iterated over all subsets averaging the results over all iterations. The broken stick model generates the outlier gene proportions, which would occur by chance alone, that is, the distribution of outlier gene proportions if there were no structure in the data.

We integrated the genomic data into the metagenomic cluster database using the “DB-update” module of the workflow. This module uses the *clusterupdate* module of MMseqs2 (Steinegger and Soding, 2017), with the same parameters used for the metagenomic clustering.

### Quality-screening of gene clusters

We examined the GCs to ensure their high intra-cluster homogeneity. We applied two methodologies to validate their cluster sequence composition and functional annotation homogeneity. We identified non-homologous sequences inside each cluster combining the identification of a new cluster representative sequence via a sequence similarity network (SSN) analysis, and the investigation of intra-cluster multiple sequence alignments (MSAs), given the new representative. Initially, we generated an SSN for each cluster, using the semi-global alignment methods implemented in *PARASAIL* (Daily, 2016) (version 2.1.5). We trimmed the SSN using a filtering algorithm (Chafee et al., 2018; Žure et al., 2017) that removes edges while maintaining the network structural integrity and obtaining the smallest connected graph formed by a single component. Finally, the new cluster representative was identified as the most central node of the trimmed SSN by the eigenvector centrality algorithm, as implemented in igraph (Csardi and Nepusz, 2006). After this step, we built a multiple sequence alignment for each cluster using *FAMSA* (Deorowicz et al., 2016) (version 1.1). Then, we screened each cluster-MSA for non-homologous sequences to the new cluster representative. Owing to computational limitations, we used two different approaches to evaluate the cluster-MSAs. We used *LEON-BIS* (Vanhoutreve et al., 2016) for the clusters with a size ranging from 10 to 1,000 genes and OD-SEQ (Jehl et al., 2015) for the clusters with more than 1,000 genes. In the end, we applied a broken-stick model (Macarthur, 1957) to determine the threshold to discard a cluster.

The predicted genes can have multi-domain annotations in different orders, therefore to validate the consistency of intra-cluster Pfam annotations, we applied a combination of w-shingling (Broder, 1997) and Jaccard similarity. We used w-shingling (k-shingle = 2) to group consecutive domain annotations as a single object. We measured the homogeneity of the *shingle sets* (sets of domains) between genes using the Jaccard similarity and reported the median similarity value for each cluster. Moreover, we took into consideration the Clan membership of the Pfam domains and that a gene might contain N-, C- and M-terminal domains for the functional homogeneity validation. We discarded clusters with a median similarity < 1.

After the validation, we refined the gene cluster database removing the clusters identified to be discarded and the clusters containing ≥ 30% *shadow genes*. Lastly, we removed the single shadow, spurious and non-homologous genes from the remaining clusters (Supp. Note 2).

### Remote homology classification of gene clusters

To partition the validated GCs into the four main categories, we processed the set of GCs containing Pfam annotated genes and the set of not annotated GCs separately. For the annotated GCs, we inferred a consensus protein domain architecture (DA) (an ordered combination of protein domains) for each annotated gene cluster. To identify each gene cluster consensus DA, we created directed acyclic graphs connecting the Pfam domains based on their topological order on the genes using *igraph* (Csardi and Nepusz, 2006). We collapsed the repetitions of the same domain. Then we used the gene completeness as a positive-weighting value for the selection of the cluster consensus DA. Within this step, we divided the GCs into “Knowns” (Known) if annotated to at least one Pfam domains of known function (DKFs) and “Genomic unknowns” (GU) if annotated entirely to Pfam domains of unknown function (DUFs). We aligned the sequences of the non-annotated GCs with FAMSA (Deorowicz et al., 2016) and obtained cluster consensus sequences with the *hhconsensus* program from *HH-SUITE* (Steinegger et al., 2019a). We used the cluster consensus sequences to perform a nested search against the UniRef90 database (release 2017_11) (The UniProt Consortium, 2017) and NCBI *nr* database (release 2017_12) (NCBI Resource Coordinators, 2018) to retrieve non-Pfam annotations with *MMSeqs2* (Steinegger and Soding, 2017) (“*-e 1e-05 --cov-mode 2 -c 0*.*6”*). We kept the hits within 60% of the Log(best-e-value) and searched the annotations for any of the terms commonly used to define proteins of unknown function (Supp. Table 12). We used a quorum majority voting approach to decide if a gene cluster would be classified as *Genomic Unknown* or *Known without Pfams* based on the annotations retrieved. We searched the consensus sequences without any homologs in the UniRef90 database against NCBI *nr*. We applied the same approach and criteria described for the first search. Ultimately, we classified as *Environmental Unknown* those GCs whose consensus sequences did not align with any of the NCBI *nr* entries.

In addition, we developed some conservative measures to control the trade-off between specificity and sensitivity for the remote homology searches such as (1) a modification of the algorithm described in Hingamp et al. (Hingamp et al., 2013) to get a confident group of homologs to determine if a query protein is known or unknown by a quorum majority voting approach (Supp. Note 3); (2) strict parameters in terms of iterations, bidirectional coverage and probability thresholds for the HHblits alignments to minimize the inclusion of non-homologous sequences; and (3) avoid providing annotations for our gene clusters, as we believe that annotation should be a careful process done on a smaller scale and with experimental context.

### Gene cluster remote homology refinement

We refined the *Environmental Unknown* GCs to ensure the lack of any characterization by searching for remote homologies in the Uniclust database (release 30_2017_10) using the HMM/HMM alignment method *HHblits* (Remmert et al., 2012). We created the HMM profiles with the *hhmake* program from the *HH-SUITE* (Steinegger et al., 2019a). We only accepted those hits with an *HHblits-probability* ≥ 90% and we re-classified them following the same majority vote approach as previously described. The clusters with no hits remained as the refined set of EUs. We applied a similar refinement approach to the KWP clusters to identify GCs with remote homologies to Pfam protein domains. The KWP HMM profiles were searched against the Pfam *HH-SUITE* database (version 31), using *HHblits*. We accepted hits with a probability ≥ 90% and a target coverage > 60% and removed overlapping domains as described earlier. We moved the KWP with remote homologies to known Pfams to the Known set, and those showing remote homologies to Pfam DUFs to the GUs. The clusters with no hits remained as the refined set of KWP.

### Gene cluster characterization

We used the *MMseqs2 taxonomy* module (commit: b43de8b7559a3b45c8e5e9e02cb3023dd339231a) in combination with the UniProtKB (release of January 2018) (UniProt Consortium, 2018) to retrieve the taxonomic ids of all genes in a gene cluster. The *taxonomy* module implements the 2bLCA (Hingamp et al., 2013) to compute the lowest common ancestor of query sequence. We used the following parameters “*-e 1e-05 –cov-mode 0 -c 0*.*6”* for the search. To retrieve the taxonomic lineages, we used the R package *CHNOSZ* (Dick, 2008).

We used eggNOG-mapper (Huerta-Cepas et al., 2017) and the EggNog5 database (Huerta-Cepas et al., 2019) to provide functional annotations for each gene in a gene cluster. We refined the functional annotations by selecting the orthologous group within the lowest taxonomic level predicted by EggNog-mapper.

We measured the intra-cluster taxonomic and functional admixture by applying the *entropy*.*empirical()* function from the *entropy* R package (Hausser and Strimmer, 2008). This function estimates the Shannon entropy based on the different taxonomic and functional annotation frequencies. For each cluster, we also retrieved the cluster consensus taxonomic and functional annotation using a quorum majority voting approach..

In addition to the taxonomic and functional annotations, we evaluated the clusters’ level of darkness and disorder using the Dark Proteome Database (DPD) (Perdigão et al., 2017) as reference. We searched the cluster genes against the DPD, applying the MMseqs2 search program (Steinegger and Soding, 2017) with “*-e 1e-20 --cov-mode 0 -c 0*.*6*”. For each cluster, we then retrieved the mean and the median level of darkness, based on the gene DPD annotations.

### High-quality clusters

We defined a subset of high-quality clusters based on the completeness of the cluster genes and their representatives. We identified the minimum required percentage of complete genes per cluster by a broken-stick model (Macarthur, 1957) applied to the percentage distribution. Then, we selected the GCs found above the threshold and with a complete representative.

### A set of non-redundant domain architectures

We estimated the number of potential domain architectures present in the *Known* GCs considering the large proportion of fragmented genes in the metagenomic dataset and that could inflate the number of potential domain architectures. To identify fragments of larger domain architecture, we considered their topological order in the genes. To reduce the number of comparisons, we calculated the pairwise string cosine distance (q-gram = 3) between domain architectures and discarded the pairs that were too divergent (cosine distance ≥ 0.9). We collapsed a fragmented domain architecture to the larger one when it contained less than 75% of complete genes.

### Inference of gene cluster communities

We aggregated distant homologous GCs into GCCs. The community inference approach combined an all-vs-all HMM gene cluster comparison with Markov Cluster Algorithm (MCL) (van Dongen and Abreu-Goodger, 2012) community identification. We started performing the inference on the Known GCs to use the Pfam DAs as constraints. We aligned the gene cluster HMMs using HHblits (Remmert et al., 2012) (-n 2 -Z 10000000 -B 10000000 -e 1) and we built a homology graph using the cluster pairs with probability ≥ 50% and bidirectional coverage > 60%. We used the ratio between HHblits-bitscore and aligned-columns as the edge weights (Supp. Note 9). We used MCL (van Dongen and Abreu-Goodger, 2012) (v. 12-068) to identify the communities present in the graph. We developed an iterative method to determine the optimal MCL inflation parameter that tries to maximize the relationship of five intra-/inter-community properties: (1) the proportion of MCL communities with one single DA, based on the consensus DAs of the cluster members; (2) the ratio of MCL communities with more than one cluster; (3) the proportion of MCL communities with a PFAM clan entropy equal to 0; (4) the intra-community HHblits-score/Aligned-columns score (normalized by the maximum value); and (5) the number of MCL communities, which should, in the end, reflect the number of non-redundant DAs. We iterated through values ranging from 1.2 to 3.0, with incremental steps of 0.1. During the inference process, some of the GCs became orphans in the graph. We applied a three-step approach to assigning a community membership to these GCs. First, we used less stringent conditions (probability ≥ 50% and coverage >= 40%) to find homologs in the already existing GCCs. Then, we ran a second iteration to find secondary relationships between the newly assigned GCs and the missing ones. Lastly, we created new communities with the remaining GCs. We repeated the whole process with the other categories (KWP, GU and EU), applying the optimal inflation value found for the Known (2.2 for metagenomic and 2.5 for genomic data).

### Validation of gene cluster communities

We tested the biological significance of the GCCs using the phylogeny of proteorhodopsin (Boeuf et al., 2015) (PR). We used the proteorhodopsin HMM profiles (Olson et al., 2018) to screen the marine metagenomic datasets using *hmmsearch* (version 3.1b2) (Finn et al., 2011). We kept the hits with a coverage > 0.4 and e-value <= 1e-5. We removed identical duplicates from the sequences assigned to PR with CD-HIT (Li and Godzik, 2006) (v4.6) and cleaned from sequences with less than 100 amino acids. To place the identified PR sequences into the MicRhode (Boeuf et al., 2015) PR tree first, we optimized the initial tree parameters and branch lengths with RAxML (v8.2.12) (Stamatakis, 2014). We used PaPaRA (v2.5) (Berger and Stamatakis, 2012) to incrementally align the query PR sequences against the MicRhode PR reference alignment and *pplacer* (Matsen et al., 2010) (v1.1.alpha19-0-g807f6f3) to place the sequences into the tree. Finally, we assigned the query PR sequences to the MicRhode PR Superclusters based on the phylogenetic placement. We further investigated the GCs annotated as viral (196 genes, 14 GC) comparing them to the six newly discovered viral PRs (Needham et al., 2019) using Parasail (Daily, 2016) (-a sg_stats_scan_sse2_128_16 -t 8 -c 1 -x). As an additional evaluation, we investigated the distributions of standard GCCs and HQ GCCs within ribosomal protein families. We obtained the ribosomal proteins used for the analysis combining the set of 16 ribosomal proteins from Méheust et al. (Méheust et al., 2019) and those contained in the collection of bacterial single-copy genes of anvi’o (Eren et al., 2021). Also, for the ribosomal proteins, we compared the outcome of our method to the one proposed by Méheust et al. (Méheust et al., 2019) (Supp. Note 9).

### Metagenomic sample selection for downstream analyses

For the subsequent ecological analyses, we selected those metagenomes with a number of genes larger or equal to the first quartile of the distribution of all the metagenomic gene counts. (Supp. Table 13).

### Gene cluster abundance profiles in genomes and metagenomes

We estimated abundance profiles for the metagenomic cluster categories using the read coverage to each predicted gene as a proxy for abundance. We calculated the coverage by mapping the reads against the assembly contigs using the *bwa-mem* algorithm from *BWA mapper* (Li and Durbin, 2010). Then, we used *BEDTOOLS* (Quinlan and Hall, 2010), to find the intersection of the gene coordinates to the assemblies, and normalize the per-base coverage by the length of the gene. We calculated the cluster abundance in a sample as the sum of the cluster gene abundances in that sample, and the cluster category abundance in a sample as the sum of the cluster abundances. We obtained the proportions of the different gene cluster categories applying a total-sum-scaling normalization. For the genomic abundance profiles, we used the number of genes in the genomes and normalized by the total gene counts per genome.

### Rate of genomic and metagenomic gene clusters accumulation

We calculated the cumulative number of known and unknown GCs as a function of the number of metagenomes and genomes. For each metagenome count, we generated 1000 random sets, and we calculated the number of GCs and GCCs recovered. For this analysis, we used 1,246 HMP metagenomes and 358 marine metagenomes (242 from TARA and 116 from Malaspina). We repeated the same procedure for the genomic dataset. We removed the singletons from the metagenomic dataset with an abundance smaller than the mode abundance of the singletons that got reclassified as good-quality clusters after integrating the GTDB data to minimize the impact of potential spurious singletons. To complement those analyses, we evaluated the coverage of our dataset by searching seven different state-of-the-art databases against our set of metagenomic GC HMM profiles (Supp. Note 11).

### Occurrence of gene clusters in the environment

We used 1,264 metagenomes from the TARA Oceans, MALASPINA Expedition, OSD2014 and HMP-I/II to explore the properties of the unknown sequence space in the environment. We applied the Levins Niche Breadth (NB) index (Levins, 1966) to investigate the GCs and GCCs environmental distributions. We removed the GCs and cluster communities with a mean relative abundance < 1e-5. We followed a divide-and-conquer strategy to avoid the computational burden of generating the null-models to test the significance of the distributions owing to the large number of metagenomes and GCs. First, we grouped similar samples based on the gene cluster content using the Bray-Curtis dissimilarity (Bray et al., 1957) in combination with the *Dynamic Tree Cut* (Langfelder et al., 2008) R package. We created 100 random datasets picking up one random sample from each group. For each of the 100 random datasets, we created 100 random abundance matrices using the *nullmodel* function of the *quasiswap* count method (Miklós and Podani, 2004). Then we calculated the *observed* NB and obtained the 2.5% and 97.5% quantiles based on the randomized sets. We compared the observed and quantile values for each gene cluster and defined it to have a *Narrow distribution* when the *observed* was smaller than the 2.5% quantile and to have a *Broad distribution* when it was larger than the 97.5% quantile. Otherwise, we classified the cluster as *Non-significant* (Salazar et al., 2015).

We used a majority voting approach to get a consensus distribution classification based on the ten random datasets.

### Identification of prophages in genomic sequences

We used PhageBoost (Sirén et al., 2021) to find gene regions in the microbial genomes that result in high viral signals against the overall genome signal. We set the following thresholds to consider a region prophage: minimum of 10 genes, maximum 5 gaps, single-gene probability threshold 0.9. We further smoothed the predictions using Parzen rolling windows of 20 periods and looked at the smoothed probability distribution across the genome. We disregarded regions that had a summed smoothed probability less than 0.5, and those regions that did differ from the overall population of the genes in a genome by using Kruskal–Wallis rank test (p-value 0.001).

### Lineage-specific gene clusters

We used the F1-score developed for AnnoTree (Mendler et al., 2019) to identify the lineage-specific GCs and to which rank they are specific. Following similar criteria to the ones used in Mendler et al. (Mendler et al., 2019), we considered a gene cluster to be lineage-specific if it is present in less than half of all genomes and at least 2 with F1-score > 0.95.

### Phylogenetic conservation of gene clusters

We calculated the phylogenetic conservation (τD) of each gene cluster using the *consenTRAIT* (Martiny et al., 2013) function implemented in the R package *castor* (Martiny et al., 2013). We used a paired Wilcoxon rank-sum test to compare the average τD values for lineage-specific and non-specific GCs.

### Evaluation of the OM-RGC v2 uncharacterized fraction

We integrated the 46,775,154 genes from the second version of the TARA Ocean Microbial Reference Gene Catalog (OM-RGC v2) (Salazar et al., 2019) into our cluster database using the same procedure as for the genomic data. We evaluated the uncharacterized fraction and the genes classified into the eggNOG (Huerta-Cepas et al., 2019) category S within the context of our database.

### Augmenting RB-TnSeq experimental data

We searched the 37,684 genes of unknown function associated with mutant phenotypes from Price et al. (Price et al., 2018) against our gene cluster profiles. We kept the hits with e-value ≤ 1e-20 and a query coverage > 60%. Then we filtered the results to keep the hits within 90% of the Log(best-e-value), and we used a majority vote function to retrieve the consensus category for each hit. Lastly, we selected the best-hits based on the smallest e-value and the largest query and target coverage values. We used the fitness values from the RB-TnSeq experiments from Price et al. to identify genes of unknown function that are important for fitness under certain experimental conditions.

## Availability of data and materials

The code used for the analyses in the manuscript is available at https://github.com/functional-dark-side/functional-dark-side.github.io/tree/master/scripts. The code to recreate the figures is available at https://github.com/functional-dark-side/vanni_et_al-figures. Detailed descriptions of the different methods and results of this manuscript are available at https://dark.metagenomics.eu. The workflow AGNOSTOS is available at https://github.com/functional-dark-side/agnostos-wf (the version used for the manuscript can be found here https://doi.org/10.5281/zenodo.4557847) and its database can be downloaded from https://doi.org/10.6084/m9.figshare.12459056.

## Acknowledgements

The authors thankfully acknowledge the computer resources at MareNostrum and the technical support provided by Barcelona Supercomputing Center (RES-AECT-2014-2-0085), the BMBF-funded de.NBI Cloud within the German Network for Bioinformatics Infrastructure (de.NBI) (031A537B, 031A533A, 031A538A, 031A533B, 031A535A, 031A537C, 031A534A, 031A532B), the University of Oxford Advanced Research Computing (http://dx.doi.org/10.5281/zenodo.22558) and the MARBITS bioinformatics core at ICM-CSIC. CV was supported by the Max Planck Society. AFG received funding from the European Union’s Horizon 2020 research and innovation program Blue Growth: Unlocking the potential of Seas and Oceans under grant agreement no. 634486 (project acronym INMARE). AM was supported by the Biotechnology and Biological Sciences Research Council [BB/M011755/1, BB/R015228/1] and RDF by the European Molecular Biology Laboratory core funds. EOC was supported by project INTERACTOMA RTI2018-101205-B-I00 from the Spanish Agency of Science MICIU/AEI. SGA and PS received additional funding by the project MAGGY (CTM2017-87736-R) from the Spanish Ministry of Economy and Competitiveness. The Malaspina 2010 Expedition was supported by the Spanish Ministry of Economy and Competitiveness (MINECO) through the Consolider-Ingenio program (ref. CSD2008-00077). The authors thank Johannes Söding and Alex Bateman for helpful discussions.

## Contributions

CV, MSS and AF-G performed the analyses and wrote the computational workflow. MS assisted with the clustering and remote homology searches. KS helped with the identification of prophages in genomic sequences. PLB and AB provided feedback and assisted with the ecological analyses. RDF and AM provided feedback and information on the MGnify and Pfam databases. CMD, PS and SGA provided the Malaspina metagenomes. TOD and AME analyzed data in the context of metagenome-assembled genomes. AF-G conceived the study and supervised the work. CV and AF-G wrote the manuscript. All authors read, edited and approved the final manuscript.

