## Supplementary file for "Unifying the known and unknown microbial coding sequence space"

### Supplementary figures

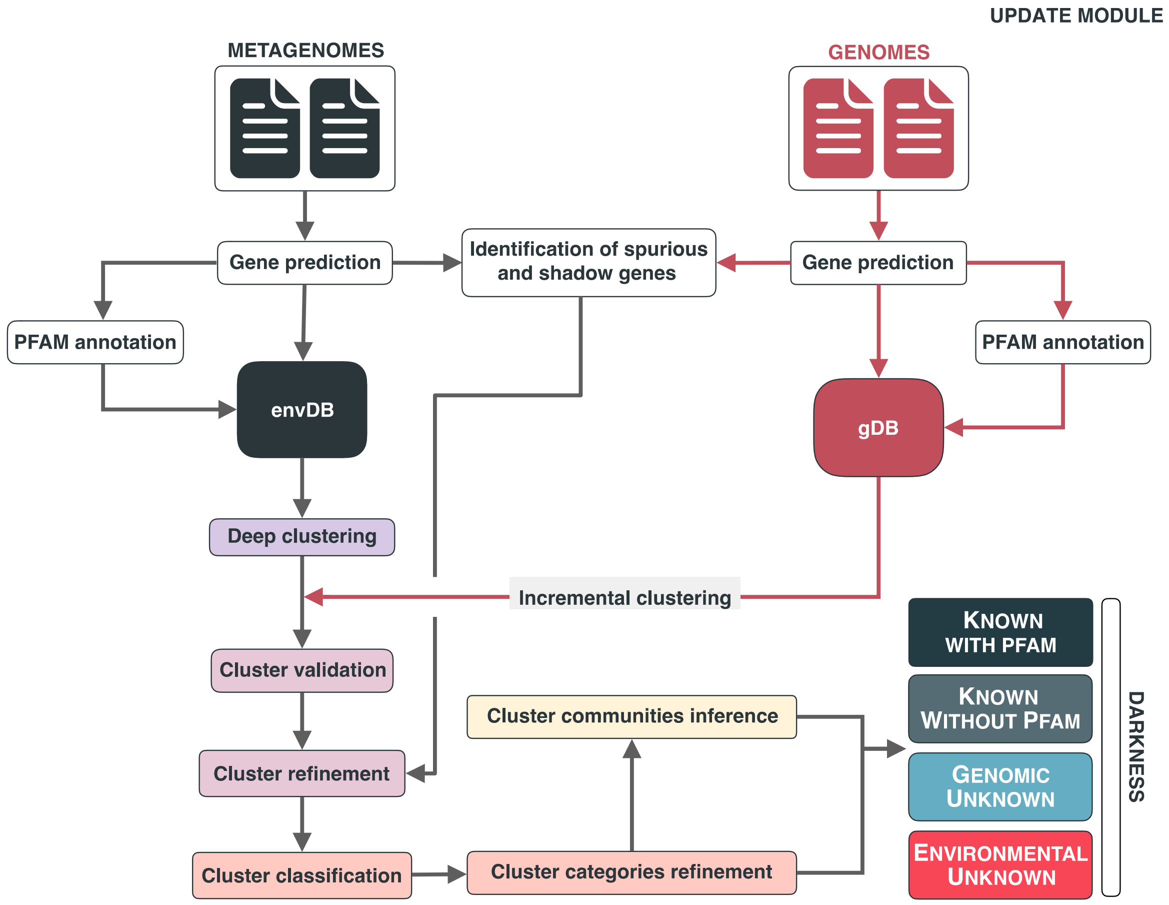


**Supplementary Figure 1.** Overview of the workflow to partition the genomic and metagenomic sequence space between known and unknown. The workflow performs gene prediction, gene clustering, gene clustering validation and refinement, GCC inference, and partitions the sequence space in the different known and unknown categories.

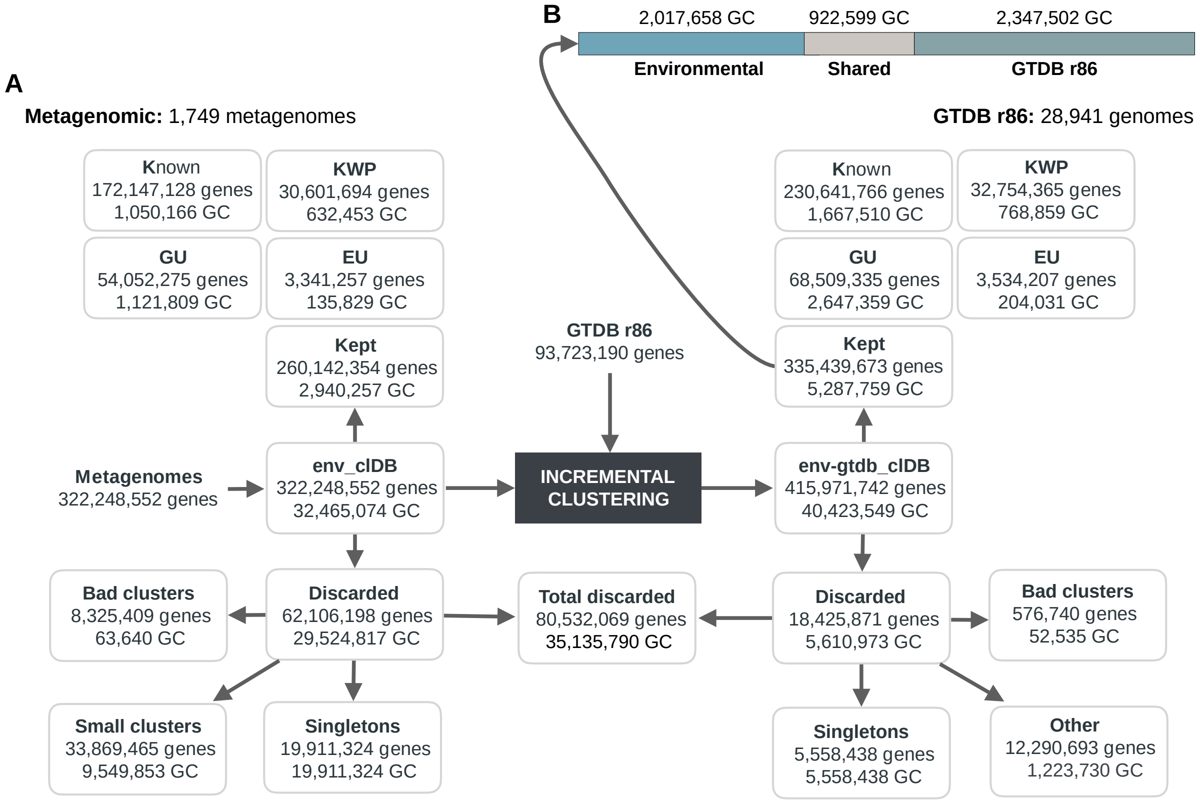


**Supplementary Figure 2.** The diagram shows a schematic description of the number of genes and GCs that have been kept or discarded. (A) We analyzed a dataset of 1,749 metagenomes from marine and human environments and 28,941 genomes from the GTDB_r86 summing up to 415,971,742 genes. The composition of the genomic box "Other" is described in supplementary Note 5. (B) GC overlap between the environmental and genomic datasets.

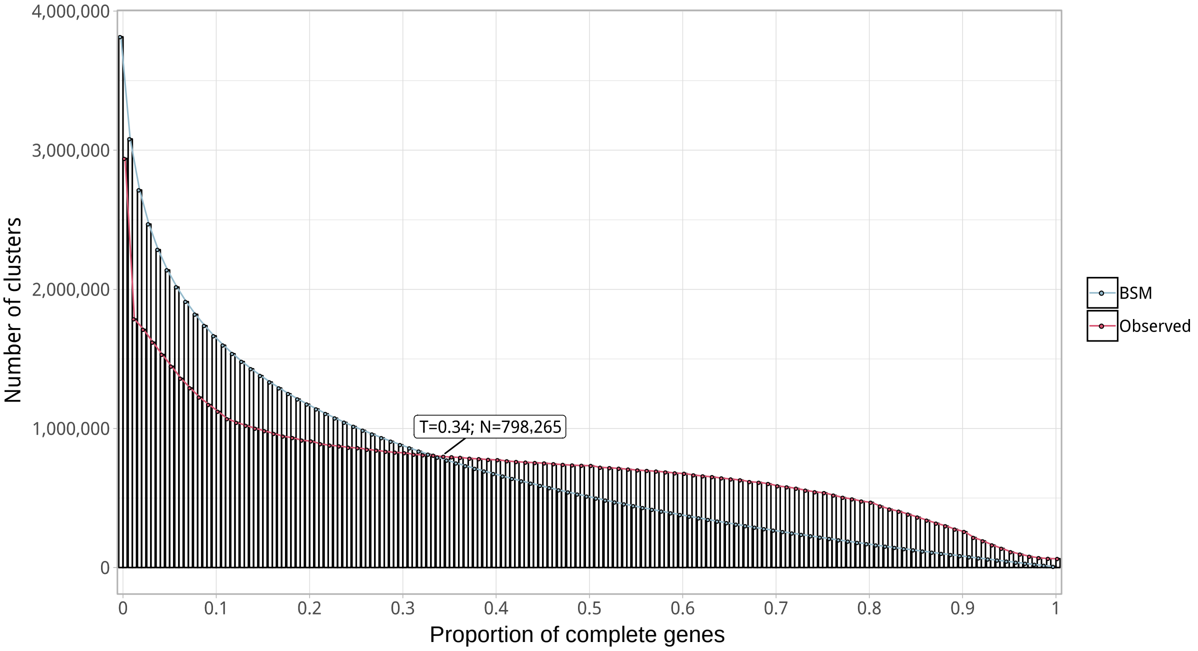


**Supplementary Figure 3.** Proportion of complete genes per cluster. Distribution of observed values compared with those generated by the Broken-stick model. The cut-off was determined at 34% complete genes per cluster.

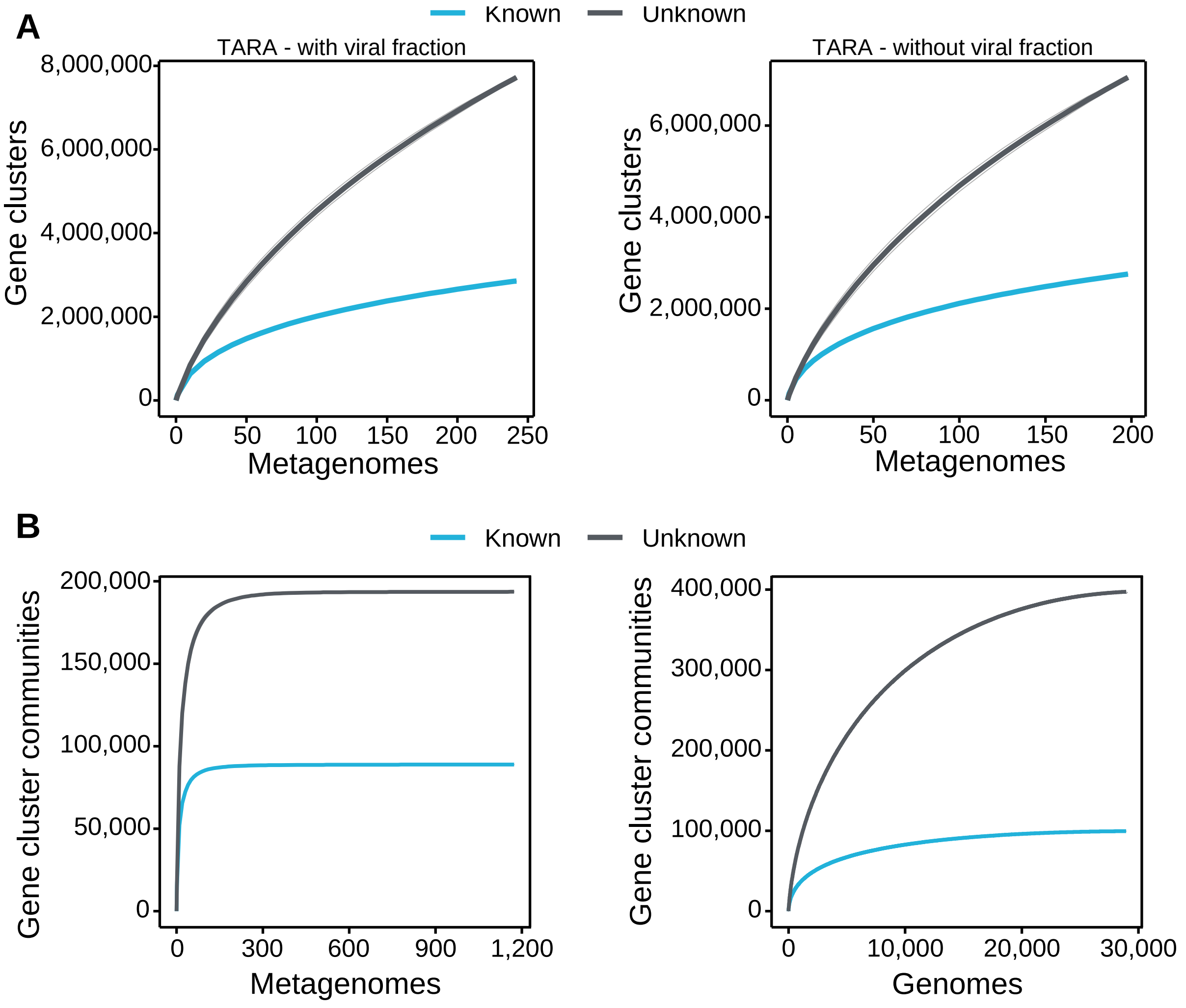


**Supplementary Figure 4:** Collector curves for the known and unknown sequence space. (A) Collector curves at the gene cluster level, for the TARA metagenomes, including the viral fraction (left) and excluding it (right) from the analysis. (B) Collector curves at gene cluster community level for the metagenomes from TARA, MALASPINA, and HMP-I/II projects (left) and the 28,941 GTDB genomes (right).

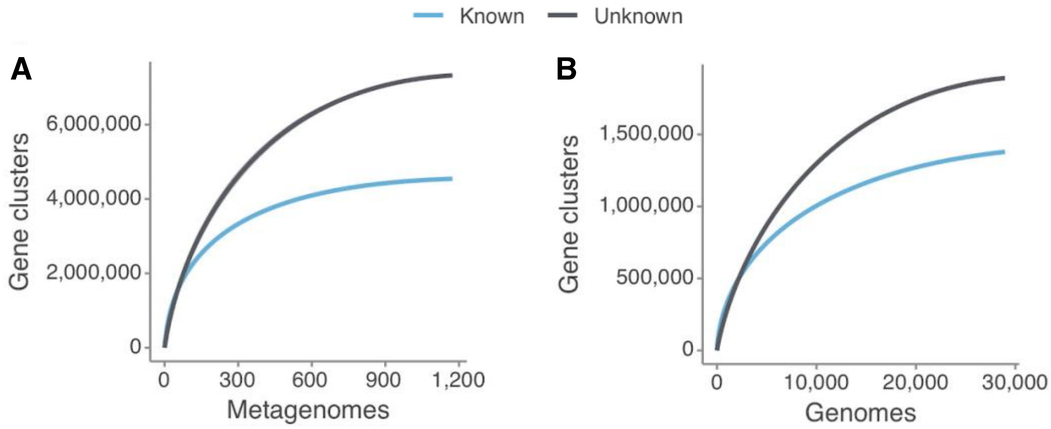


**Supplementary Figure 5:** Collector curves for the known and unknown sequence space at the gene cluster communities level for (A) the metagenomes from TARA, MALASPINA and HMP-I/II projects, and for (B) the 28,941 GTDB genomes. Singletons were excluded from the calculations.

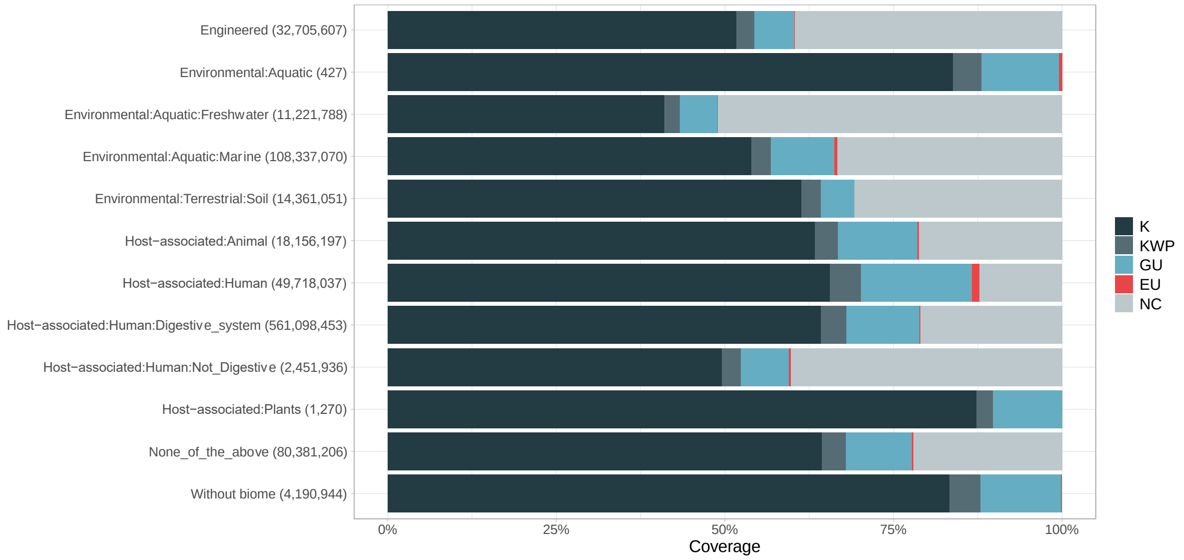


**Supplementary Figure 6.** Proportion of gene cluster categories per biome. On the y-axis are reported the 11 main biome categories indicated by MGnify and in parenthesis the total number of genes in each biome. The gray fraction represents the pool of genes from MGnify that were not found in our dataset.

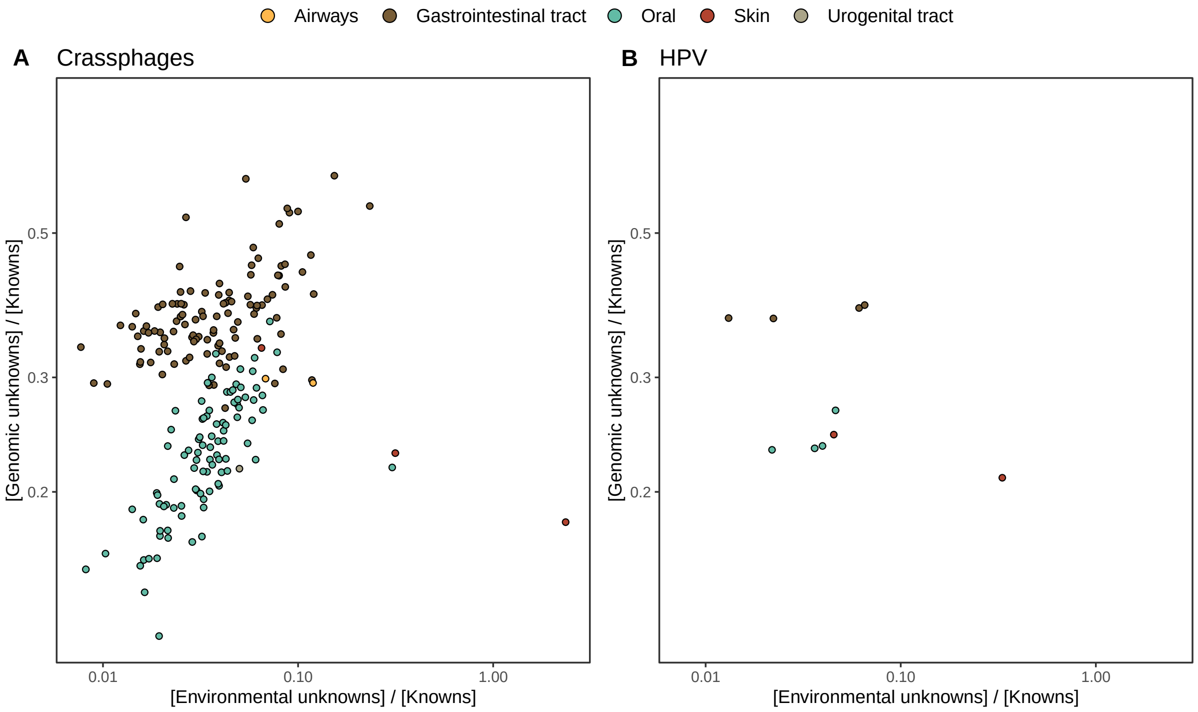


**Supplementary Figure 7.** HMP outlier samples enriched in (A) crAssphages, and (B) papillomaviruses (HPV).

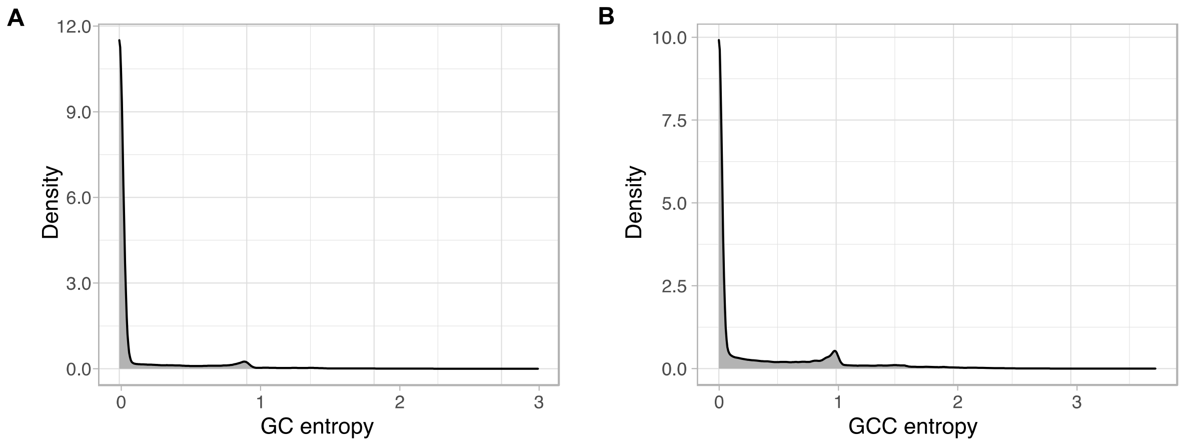


**Supplementary figure 8.** EggNOG annotations entropy within the GCs (A) and the GCCs (B)**.** The entropy was calculated using the function *entropy.empirical()* from the R package “entropy”, which estimates the Shannon entropy values based on the value empirical frequencies.

### Supplementary Tables

**Supplementary Table 1.** Number of metagenomic clusters and genes after the validation and refinement steps.

| | Good-quality | Bad-quality | Total |
|---|---|---|---|
| Clusters | 2,940,257 | 63,640 | 32,465,074 |
| Genes | 260,142,354 | 8,325,409 | 322,248,552 |

**Supplementary Table 2.** MG + GTDB high quality (HQ) subset of gene clusters (GCs).

| Category | HQ GCs | HQ genes | pHQ GCs | pHQ genes |
|---|---|---|---|---|
| K | 76,718 | 40,710,936 | 0.0145 | 0.120 |
| KWP | 16,922 | 1,733,599 | 0.00320 | 0.005132 |
| GU | 95,370 | 9,908,630 | 0.0180 | 0.0293 |
| EU | 14,207 | 477,625 | 0.00269 | 0.00141 |
| Total | 203,217 | 52,830,790 | 0.0384 | 0.1562 |

**Supplementary Table 3.** Mean proportion of complete genes per cluster in the four functional categories.

| | K | KWP | GU | EU |
|---|---|---|---|---|
| Mean percentage of complete genes | 0.50 | 0.22 | 0.68 | 0.70 |

**Supplementary Table 4.** KWP high-quality gene clusters (GCs) distribution in the COG groups. (Full table in Supplementary_tables_1.xlsx)

| COG group | Number of GCs | Proportion of GCs |
| --- | --- | --- |
| CELLULAR PROCESSES AND SIGNALING | 2,292 | 0.135 |
| INFORMATION STORAGE AND PROCESSING | 1,582 | 0.0935 |
| METABOLISM | 1,679 | 0.0992 |
| POORLY CHARACTERIZED | 2,899 | 0.171 |
| NC | 8,470 | 0.501 |

**Supplementary Table 5.** MG + GTDB gene clusters summary statistics. (Supplementary_tables_2.xlsx)

**Supplementary Table 6.** Environmental (metagenomic) dataset description.

(A) Number of samples and sites per metagenomic project.

| Dataset | Reference | Samples | Sites | Contigs |
|---|---|---|---|---|
| TARA | Sunagawa et al. (Sunagawa et al., 2015) | 242 | 141 | 62,404,654 |
| Malaspina | Duarte et al. (Duarte, 2015) | 116 | 30 | 9,330,293 |
| OSD | Kopf et al. (Kopf et al., 2015) | 145 | 139 | 4,127,095 |
| HMP | Lloyd-Price et al. (Lloyd-Price et al., 2017) | 1,246 | 18 | 80,560,927 |

| Dataset | Reference | Samples | Sites | Reads |
|---|---|---|---|---|
| GOS | Rusch et al. (Rusch et al., 2007) | 80 | 70 | 12,672,518 |

(B) Number of predicted genes per completeness category.

| Total | "00" | "10" | "01" | "11" |
|---|---|---|---|---|
| 322,248,552 | 118,717,690 | 106,031,163 | 102,966,482 | 75,694,123 |

Note: "00"=complete, both start and stop codon identified. "01"=right boundary incomplete. "10"=left boundary incomplete. "11"=both left and right edges incomplete.

**Supplementary Table 7.** Proportion of genes in each cluster category, and Pfam amino acids coverage per cluster category. (Supplementary_tables_1.xlsx)

**Supplementary Table 8.** List of HMP outlier samples (Supplementary_tables_1.xlsx).

**Supplementary Table 9.** Summary of the number of EU clusters based on their presence in MAGs and their environmental distribution, obtained with the Levin's Niche Breadth index.

| | Total clusters | Broad | Narrow | Non-significant |
|---|---|---|---|---|
| Total EU | 204,031 | 471 | 8,421 | 195,079 |
| EU in MAGs | 55,520 | 88 | 316 | 55,116 |
| EU not in MAGs | 148,511 (73%) | 383 (81%) | 8,105 (96%) | 140,023 (72%) |

**Supplementary Table 10.** Number of phylogenetic conserved and lineage-specific gene clusters (GCs) in the GTDB bacterial phylogeny. (Supplementary_tables_1.xlsx).

**Supplementary Table 11.** Clusters in the GU community GU_c_21103 (Supplementary_tables_1.xlsx).

**Supplementary Table 12.** Number of lineage-specific gene clusters of unknown function at different taxonomic levels within the *Cand. Patescibacteria* phylum.

| Taxonomic level | Number of clusters |
|---|---|
| Phylum | 2 |
| Class | 6 |
| Order | 104 |
| Family | 1,456 |
| Genus | 6,987 |
| Species | 45,788 |

**Supplementary Table 13.** List of filtered samples used for the metagenomic analyses. (Supplementary_tables_1.xlsx)

**Supplementary Table 14.** List of terms commonly used to define proteins of unknown function in public databases. (Supplementary_tables_1.xlsx)

**Supplementary Table 15.** Shannon entropy values for the eggNOG annotations within the gene clusters.

| | Min. | 1st Qu. | Median | Mean | 3rd Qu. | Max. |
|---|---|---|---|---|---|---|
| Entropy per GC | 0.000 | 0.000 | 0.000 | 0.105 | 0.000 | 3.729 |

**Supplementary Table 16.** Shannon entropy values for the eggNOG annotations within the gene clusters communities.

| | Min. | 1st Qu. | Median | Mean | 3rd Qu. | Max. |
|---|---|---|---|---|---|---|
| Entropy per GCC | 0.000 | 0.000 | 0.000 | 0.285 | 0.400 | 3.721 |

### Supplementary Notes

#### Supplementary Note 1 - Metagenomic singletons and small gene clusters

*Analysis of metagenomic singletons and gene clusters with less than ten genes.*

The singletons represent 60% of the gene clusters (GCs) and 6% of the total genes. The GCs with less than ten genes, here referred to as small GCs for simplicity, represent 29% of the GCs and 10.5% of the gene dataset (Supp. Fig. 2A). Although we discarded these two sets from the main study, we investigated them to obtain a complete analysis of the initial dataset. Both sets were first searched against the Pfam database of protein domain families (Finn et al., 2016), and subsequently classified following the steps described in Supplementary Note 3. For the small GCs classification, we used the cluster consensus sequence, which we extracted using the *hhconsensus* program of the HH-SUITE (Steinegger et al., 2019), from the GC multiple sequence alignments (MSAs), generated with FAMSA (Deorowicz et al., 2016).

We could not find any homologous in the Pfam database for the large majority of both singletons and small GCs, 95%, and 89%, respectively (Supp. Table 1-1). After the classification, the large majority of the singletons remained completely uncharacterized, (64% was identified as EU) (Supp. Table 1-2). Similarly, the small GCs were also found dominated by GCs of unknowns, with 38% of the clusters classified as EU and 29% as GU (Supp. Table 1-2).

**Supplementary Table 1-1.** Singletons and small GCs Pfam annotations.

| | Total | Annotated | Not annotated |
|---|---|---|---|
| Singletons | 19,911,324 | 934,548 | 18,976,776 |
| Small GCs | 9,549,853 | 1,028,076 | 8,521,777 |

**Supplementary Table 1-2.** Number of singletons and small GCs per functional category.

| | K | KWP | GU | EU |
|---|---|---|---|---|
| Singletons | 852,413 | 3,505,161 | 2,763,476 | 12,790,274 |
| Small GCs | 946,112 | 2,213,654 | 2,744,262 | 3,645,825 |

### Supplementary Note 2 - Metagenomic gene cluster validation and refinement

*To obtain a set of gene clusters characterized by a high intra-cluster homogeneity, we identified spurious, shadow and outlier genes, and we removed them from the clusters.*

*Identification of spurious genes.* We identified spurious genes by screening our gene data set against the *AntiFam* database (Eberhardt et al., 2012).
*Identification of shadow genes.* We identified shadow genes using the procedure described in Yooseph et al. (Yooseph et al., 2008). (1) Two genes on the same strand are considered overlapping if their intervals overlap by at least 60 bps; (2) genes that are on the opposite strands are considered overlapping if their intervals overlap by at least 50 bps, and their 3' ends are within each other's intervals, or if their intervals overlap by at least 120 bps and the 5' end of one is in the interval of the other.
*Identification of outlier genes.* Outlier genes are sequences inside a cluster non-homologous to the other cluster genes and were identified during the cluster validation step (see Methods - **Gene cluster validation**).
The number of spurious, shadow and outlier genes identified in the data set is reported in Supplementary Table 2-1.
*Cluster refinement.* After the validation, we proceeded with the retrieval of the subset of "good" clusters. Clusters with ≥ 30% shadow genes were identified as shadow-clusters, as proposed in Yooseph et al. (Yooseph et al., 2008). During the cluster validation, we identified a minimum of 10% outlier genes as the threshold to classify a cluster as "bad-quality" (Supp. Fig. 2-2; Suppl. Table 2-2A). We combined this threshold with a Jaccard similarity index < 1, indicating a low intra-cluster Pfam domain architecture (DA) homogeneity, for the Pfam annotated clusters (Supp. Table 2-2B). We performed the cluster refinement in three consecutive steps:

I. Discard the "bad" clusters (≥ 10% outliers & Jaccard similarity index <1)
II. Discard the "shadow" clusters (≥ 30% shadow genes)
III. Remove the single shadow, spurious and outlier genes from the remaining clusters.

The results for each step are shown in Supplementary Table 2-3. From the initial set of ~3M clusters with more than ten genes, we identified 57,052 GCs as "bad" and 6,261 as "shadow". From the remaining set of 2,940,593 clusters, we removed a total of 2,708,994 shadow, spurious and outlier genes. During this last step, we discarded 336 more clusters: 244 resulted being composed only of spurious and outlier genes (one in the Pfam annotated set of clusters and 243 in the non-annotated set), and 92 clusters were discarded since they were left as singletons after refinement. Besides, we moved 1,190 Pfam annotated clusters to the non-annotated set since they were left without any annotated gene. In summary, we removed 63,640 GCs and a total of 8,325,409 genes, respectively, 2% and 3% of the initial data set. The refined set contains 2,940,592 GCs and 260,142,354 genes (Supp. Table 3).

**Supplementary Table 2-1.** Number of spurious, shadow and outlier genes in the metagenomic clusters.

| Gene category | Clusters ≥ 10 genes | Clusters < 10 genes | Singletons |
|---|---|---|---|
| Spurious | 44,205 | 6,784 | 2,335 |
| Shadow | 289,258 | 144,571 | 177,126 |
| Outliers | 3,118,850 | - | - |

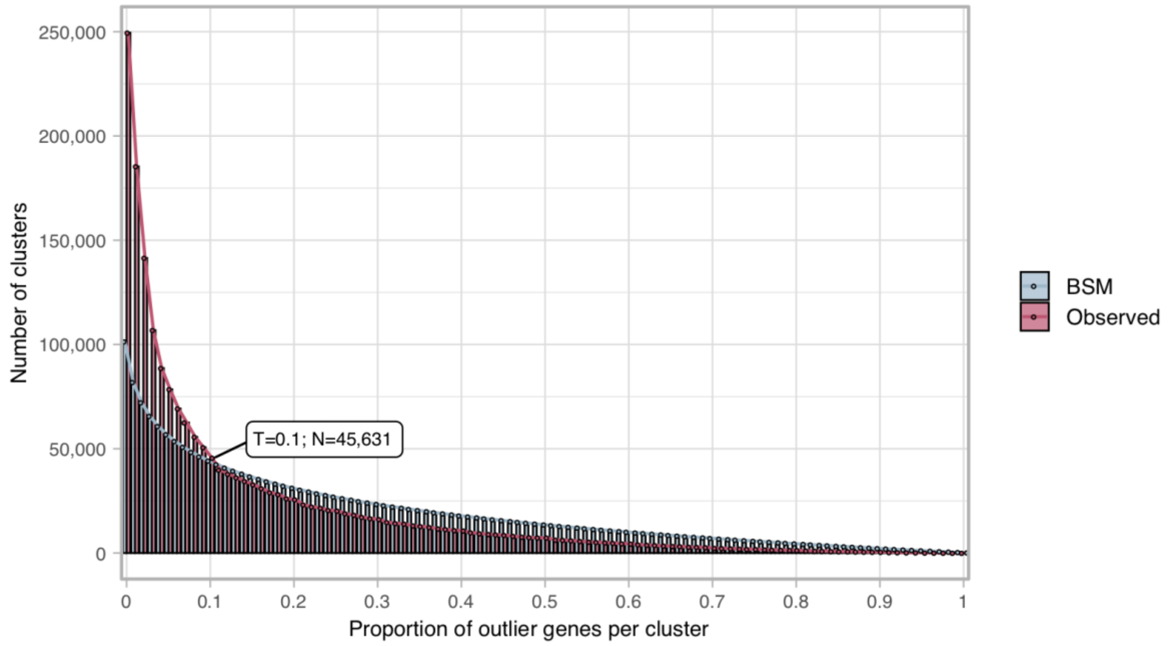


**Supplementary Figure 2-1.** Proportion of outlier genes detected within each cluster MSA. Distribution of observed values compared with those generated by the Broken-stick model. The cut-off was determined at 10% outlier genes per cluster.

**Supplementary Table 2-2.** Metagenomic gene cluster validation results.

(A) Evaluation of cluster sequence composition.

| | Pre-Compos. validation | good quality | bad quality |
|---|---|---|---|

| | | | |
|---|---|---|---|
| Clusters | 3,003,897 | 2,958,266 | 45,631 |
| Genes | 268,467,763 | 266,268,638 | 2,199,125 |

(B) Evaluation of cluster Pfam functional annotations.

| | Pre-Funct. validation | Funct. good | Funct. bad |
|---|---|---|---|
| Clusters | 1,015,924 | 1,004,166 | 11,758 |
| Genes | 181,433,541 | 178,167,583 | 3,246,002 |

**Supplementary Table 2-3**

Steps:

Step I - Removing of the "bad clusters"

Step II - Removing of the "shadow clusters"

Step III - Removing single spurious, shadow or outlier genes

(A) Number of clusters in each step of the cluster refinement.

| | Step I | Step II | Step III | Refined |
|---|---|---|---|---|
| Clusters | 3,003,897 | 2,946,845 | 2,940,593 | 2,940,257 |
| Removed | -57,052 | -6,252 | -336 | |

(B) Number of genes in each step of the cluster refinement.

| | Step I | Step II | Step III | Refined |
|---|---|---|---|---|
| Genes | 268,467,763 | 263,022,636 | 262,851,348 | 260,142,354 |
| Removed | -5,445,127 | -171,288 | -2,708,994 | |

### Supplementary Note 3 - Metagenomic gene cluster classification and remote homology refinement

*Classification of the refined subset of gene clusters and remote homology refinement.*

**Methods**

We searched the gene clusters (GCs) without any Pfam annotated gene against two functional databases, the UniRef90, from UniProt (The UniProt Consortium, 2017), and the NCBI *nr* database (NCBI Resource Coordinators, 2018). We screened the two databases using the cluster consensus sequences, obtained by applying the *hhconsensus* program of the *HH-SUITE (Steinegger et al., 2019)* on the clusters multiple sequence alignments (MSAs) generated with the *FAMSA* program (Deorowicz et al., 2016). We performed two nested searches using the *MMSeqs2 (Steinegger & Söding, 2017)* program and following a similar workflow as the "2bLCA" described in Hinghamp et al. (Hingamp et al., 2013). The search-workflow consisted of five steps: First, we searched the consensus sequences against the functional database, with -e 1e-05 --cov-mode 2 -c 0.6. Second, we extracted the high scoring pairs (HSP) of the best hits and we searched them again using the same parameters. Third, we merged the top hits from the first with the second search results. Fourth, we filtered out the second search hits with a bigger e-value than the first search top hits. And fifth, we selected the hits that were found in 60% of the log10(best-e-value). We first applied this search-workflow to screen the UniRef90 database (release 2017_11) (The UniProt Consortium, 2017). We classified the GCs as GU if their consensus sequences were found annotated to proteins labeled with any of the terms commonly used to define proteins of unknown function in public databases (Supp. Table 14). We classified, instead, as KWP, the clusters with consensus annotated to functionally characterized proteins. Secondly, we applied the same search-workflow to search the consensus sequences with no homologs in the UniRef90 database, against the NCBI *nr* database (release 2017_12) (NCBI Resource Coordinators, 2018). We used the same criteria to classify a GC as GU or KWP. Ultimately, we classified as EU the GCs whose consensus sequences did not align with any of the NCBI *nr* entries.

We processed the Pfam annotated GCs to retrieve a GC consensus domain architecture (DA). We classified as GU the GCs with a consensus DA composed only of Pfam domain of unknown function (DUFs) and as K the rest. The methods for this step are described in Methods - **Remote homology classification of gene clusters**.

We refined the classified GCs to account for remote homologies. A detailed description of this process can be found in Methods - **Gene cluster remote homology refinement**.

**Results**

From the 1,946,737 non-annotated clusters, 1,581,115 were found homologous to UniRef90 entries. Of these hits, more than 50% were found homologous to "hypothetical" proteins and classified as GU, and the other hits were labeled as KWP. The remaining 365,622 clusters, with no homologs to UniRef90, were screened against the NCBI nr database. We found 20,277 clusters in the NCBI nr, of them, 15,998 clusters were homologous to "hypothetical" proteins, and 4,279 clusters to characterized proteins and were classified respectively as GU and KWP. The remaining 345,345 clusters were not found in the NCBI nr database and therefore identified as EU. After the cascaded profile search against UniRef90 and NCBI nr, and the analysis of the GC consensus DAs, we classified the GCs into 912,551 K, 753,718 KWP, 928,643 GU, and 345,345 EU. Detailed results for each search are reported in Supplementary Table 3-1.

**Supplementary Table 3-1.** Metagenomic gene clusters classification steps.

(A) Results from the search against the UniRef90 database

| **Search vs UniRef90** | Hits | | No-hits |
|---|---|---|---|
| Initial clusters:1,946,737 | 1,581,115 | | 365,622 |
| | Characterized | Hypothetical | |
| | 749,439 | 831,676 | |

(B) Results from the search against the and the NCBI nr databases

| **Search vs NCBI nr** | Hits | | No-hits |
|---|---|---|---|
| Initial clusters: 365,622 | 20,277 | | 345,345 |
| | Characterized | Hypothetical | |
| | 4,279 | 15,998 | |

(C) Classification of the Pfam annotated GCs based on the consensus DAs.

| **Consensus DA analysis** | Annotated to DKF DAs | Annotated to DUF DAs |
|---|---|---|
| Initial clusters: 993,520 | 912,551 | 80,969 |

**Supplementary Table 3-2.** Metagenomic GC remote homology refinement steps.

| | K | KWP | GU | EU |
|---|---|---|---|---|
| Initial GCs | 912,551 | 753,718 | 928,643 | 345,345 |

| | | | | |
|---|---|---|---|---|
| EU refinement | - | +38,333 | +171,183 | -209,516 |
| Post-EU refinement | 912,551 | 792,051 | 1,099,826 | 135,829 |
| KWP refinement | +137,615 | -159,598 | +21,983 | - |
| Refined GCs | 1,050,166 | 632,453 | 1,121,809 | 135,829 |

#### Supplementary Note 4 - GTDB integration

*Results from the integration of the Genome Taxonomy Database(Parks et al., 2018) into the metagenomic dataset.*

We integrated the metagenomic GCs with the 93,723,190 genes from the archaeal and bacterial GTDB genomes (release 86) (Parks et al., 2018). A total of 67,446,376 genomic genes, 72% of the whole dataset, were found in the metagenomic GCs. The remaining 26,276,814 (28% of the initial dataset) genes were then clustered separately into 7,958,475 genomic GCs (Supp. Table 4-1). This set of GCs was processed through our workflow steps to be validated, classified and refined.

Within the set of genomic GCs, we identified 5,558,438 singletons and 2,400,037 GCs with more than one gene. We were able to annotate to Pfam protein domain families 41% of the genomic genes. The annotation led to 556,834 annotated GCs and 1,843,203 non-annotated GCs. The validation step determined the minimum proportion of outlier genes per cluster at 11% (Supp. Fig. 4-1). The majority of the genomic GCs showed high intra-cluster homogeneity, both in terms of sequence composition and functional annotations (Supp. Table 4-2).

After the validation, we refined the GCs removing the GCs identified as "bad" and the detected outliers' genes (see Supp. Table 4-3). We classified the refined subset of 2,347,502 GCs into the four functional categories via the same protocol applied for the metagenomic data set. The results of the GC classification are reported in Supplementary Table 4-4. After the classification steps, we refined the EU and KWP GCs searching their HMMs profiles for remote homologies in the Uniclust (release 30_2017_10) (Mirdita et al., 2017) and the Pfam (v. 31.0) (Finn et al., 2016) databases, respectively, using HHblits (Remmert et al., 2012). An overview of the results step-by-step can be found in Supplementary Table 4-5A. In the end, we obtained 617,344 GCs classified as Known, 136,406 as KWP, 1,525,550 as GU and 68,202 as EU (Supp. Table 4-5B). The genomic dataset appeared highly dominated by the GU, which accounts for 65% of the GCs. In the end, we retrieved a subset of genomic "High Quality" (mostly complete) GCs (Supp. Table 4-6). The numbers of genes and GCs for the integrated (MG+GTDB) dataset are reported in Supplementary Table 4-7.

**Supplementary Table 4-1.** GTDB integration in the metagenomic dataset.

| | Metagenomic | Shared | Genomic | Total |
|---|---|---|---|---|
| GCs | 30,301,693 | 2,163,381 | 7,958,475 | 40,423,549 |
| Genes | 199,693,614 | 190,001,314 | 26,276,814 | 415,971,742 |

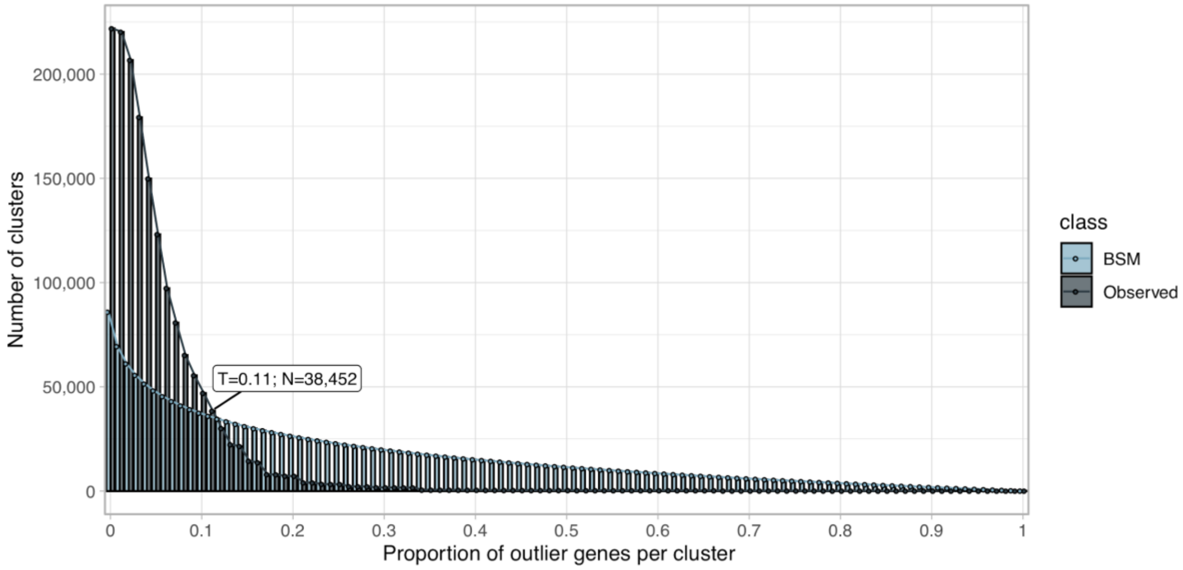


**Supplementary Figure 4-1.** Proportion of outlier genomic genes identified within each cluster MSA. Distribution of observed values compared with those of the Broken-stick model.

**Supplementary Table 4-2.** Genomic GC validation results.

(A) Evaluation of cluster sequence composition.

| | Pre-Compos. validation | good quality | bad quality |
|---|---|---|---|
| GCs | 2,400,037 | 2,361,585 | 38,452 |
| Genes | 20,718,376 | 20,364,454 | 353,922 |

(B) Evaluation of Pfam functional annotations.

| | Pre-Funct. validation | good quality | bad quality |
|---|---|---|---|
| GCs | 556,834 | 542,410 | 14,424 |
| Genes | 10,091,203 | 9,865,550 | 225,653 |

(C) Combined cluster validation results.

| | Pre-validation | good quality | bad quality |
|---|---|---|---|
| GCs | 2,400,037 | 2,347,502 | 52,535 |
| Genes | 20,718,376 | 20,141,636 | 576,740 |

**Supplementary Table 4-3.** Spurious, shadow and outlier genes in the genomic GCs.

| Gene category | GCs >= 2 genes | Singletons |
|---|---|---|
| Spurious | 3,252 | 1,312 |
| Shadow | 223,535 | 125,262 |
| Outliers | 449,080 | - |

**Supplementary Table 4-4.** Non-annotated genomic GC classification.

(A) Results from the search against the UniRef90 database.

| **Search vs UniRef90** | Hits | | No-hits |
|---|---|---|---|
| Initial GCs: 1,816,999 | 1,570,094 | | 246,905 |
| | Characterized | Hypothetical | |
| | 304,004 | 1,266,090 | |

(B) Results from the search against the NCBI nr database.

| **Search vs NCBI nr** | Hits | | No-hits |
|---|---|---|---|
| Initial GCs: 246,905 | 28,704 | | 218,201 |
| | Characterized | Hypothetical | |
| | 1,280 | 27,424 | |

(C) Classification of the Pfam annotated GCs based on the consensus DAs.

| **Consensus DA analysis** | DKF DAs | DUF DAs |
|---|---|---|
| Initial GCs: 993,520 | 912,551 | 65,688 |

**Supplementary Table 4-5.** Genomic GC remote homology refinement and final genomic GC dataset.

(A) Remote-homology refinement steps.

| | K | KWP | GU | EU |
|---|---|---|---|---|
| Initial GCs | 464,815 | 305,284 | 1,359,202 | 218,201 |
| EU refinement | - | +5,704 | +144,295 | -149,999 |
| Post-EU refinement | 464,815 | 310,988 | 1,503,497 | 68,202 |
| KWP refinement | +152,529 | -174,582 | +22,053 | - |
| Refined GCs | 617,344 | 136,406 | 1,525,550 | 68,202 |

(B) Genomic GC refined dataset.

| | K | KWP | GU | EU | Total |
|---|---|---|---|---|---|
| Genes | 9,997,529 | 663,107 | 9,305,621 | 175,379 | 20,141,636 |
| GCs | 617,344 | 136,406 | 1,525,550 | 68,202 | 2,347,502 |

**Supplementary Table 4-6.** Genomic high quality (HQ) GCs.

| Category | HQ GCs | HQ genes | pHQ GCs | pHQ genes |
|---|---|---|---|---|
| K | 12,202 | 25,105,156 | 0.0198 | 0.0096 |
| KWP | 4,019 | 1,349,165 | 0.0295 | 0.0214 |
| GU | 12,699 | 8,403,393 | 0.0083 | 0.0062 |
| EU | 438 | 471,820 | 0.0064 | 0.0074 |

**Supplementary Table 4-7.** MG + GTDB seed database. Integrated number of genes and GCs per category.

| | K | KWP | GU | EU | Total |
|---|---|---|---|---|---|
| Genes | 230,641,76 | 32,754,365 | 68,509,335 | 3,534,207 | 335,439,673 |
| GCs | 1,667,510 | 768,859 | 2,647,359 | 204,031 | 5,287,759 |

### Supplementary Note 5 – Summary of the post-genomic integration dataset

*In-detail description of the integrated metagenomic-genomic dataset.*

The integration of 93,723,190 genomic genes into the metagenomic dataset (322,248,552 genes, 32,465,074 GCs) resulted into a dataset of 415,971,742 genes and 40,423,549 GCs (Supp. Fig. 2A and Supp. table 4-1). As shown in Supp. Figure 2A, the integrated dataset is divided into: (1) “kept” GCs and (2) “discarded” GCs.

*1. The “kept” GCs.*

The “kept” GC dataset contains the 2,940,257 metagenomic “kept” GCs with 260,142,354 genes (Supp. Fig. 2A), the genomic “kept” 2,347,502 GCs with 20,141,636 genes (Supp. Table 4-5B), plus 55,155,683 genomic genes found in the metagenomic set of “kept” GCs (Supp. Table 5-1), for a total of 5,287,759 GCs and 335,439,673 genes. A description of the integrated “kept” dataset numbers of GCs and genes, and their distribution in the different categories can be found in Supp. Figure 2A and Supp. Table 4-7.

2. The “discarded” GCs.

The metagenomic “discarded” set includes 8,325,409 genes and 63,640 GCs classified as “bad” during the validation and refinement processes (Supp. Note 2), 19,911,324 singletons and 33,869,465 genes in 9,549,853 small GCs, i.e. clusters with less than 10 genes (Supp. Note 1), for a total of 62,106,198 genes and 29,524,817 GCs.

The genomic “discarded” dataset consists of 576,740 genes and 52,535 GCs classified as “bad”, 5,558,438 singletons (Supp. Note 4) and 12,290,693 genomic genes found in 1,223,730 metagenomic discarded clusters. This last set of genes, labeled as “Other” in Supp. Figure 2A, includes 1,578,862 genomic genes found in the set of metagenomic “bad” clusters, 7,010,987 genomic genes found in the metagenomic small GCs and 3,700,844 genomic genes homologous to metagenomic singletons (Supp. Table 5-1).

The integration of the metagenomic and genomic “discarded” sets resulted in 80,532,069 genes and 35,135,790 GCs.

As described above, with the integration of genomic data we enriched metagenomic singletons and small GCs. This addition resulted in a set of 52,758 metagenomic singletons and 187,953 metagenomic small GCs becoming GCs with more than ten genes. We validated and classified the 240,711 GCs in this set. We obtained 223,229 good-quality GCs, divided into 17,383 K, 89,205 KWP, 109,636 GU and 7,005 EU.

**Supplementary Table 5-1.** Overview of genomic genes found homologous to metagenomic genes.

| | Total | In MG good-quality GCs | In MG small GCs | In MG singletons | In MG bad-quality GCs |
|---|---|---|---|---|---|
| Genes | 67,446,376 | 55,155,683 | 7,010,987 | 3,700,844 | 1,578,862 |

#### Supplementary Note 6 - Gene cluster additional information

*Additional information on the metagenomic and genomic (MG + GTDB) gene cluster dataset.*

We retrieved a set of statistics for the MG + GTDB GC dataset, including the proportion of complete genes per cluster, the average gene length, the cluster level of darkness and disorder, and a cluster consensus taxonomic affiliation. The methods we applied to obtain these statistics are described in the Methods-Gene cluster characterization paragraph. Overall the K category has the largest average GC size, 139.6 genes (and a max of 168,822 genes). The average GC size is then decreasing from the known to the unknown categories, with the EU presenting the smallest average size, with 17.36 genes per GC. Similarly, the K GCs have, on average, the longest genes (258.55 aa), followed by the GU (177.16 aa), the KWP (133.22 aa) and the EU (130.65 aa). The unknown categories (GU and EU) have the highest level of completion, i.e., the proportion of complete genes per GC. The KWP GCs contain the smallest percentage of complete genes. We evaluated the levels of darkness and disorder of the GCs using the information on the DPD (Perdigão et al., 2017) annotations (Supp. Table 6-1). The categories K, KWP and GU showed a degree of darkness inversely proportional to their functional characterization. Interestingly the KWP presented the highest level of disorder (Supp. Table 6, Supp. Fig 3), while the proper characterization of these proteins is beyond the scope of this paper, our preliminary analyses suggest that KWP are enriched in intrinsically disordered proteins (Habchi et al., 2014) (Supp. Table 6-1). These proteins, usually involved in signaling and regulatory functions, don't have a well-defined 3-D structure and they can adopt many different conformations.

We used the taxonomy of 214,392,608 genes to evaluate the taxonomic variation within a GC and generated consensus taxonomic annotations for 2,630,338 GCs. The GCs taxonomic variation is low at higher taxonomic levels and it steadily increases towards Genus and Species (Supp. Table 5).

A general overview of the MG + GTDB main properties for the whole GCs dataset can be found in Supplementary Table 5 (Supplementary_tables_2.xlsx).

**Supplementary Table 6-1.** Number of MG + GTDB GCs annotated to the DPD per functional category.

| K | KWP | GU | EU |
|---|---|---|---|
| 374,555 | 8,874 | 22,135 | 0 |

### Supplementary Note 7 - Gene cluster communities

*Metagenomic and genomic gene cluster community inference detailed results.*

We aggregated the gene clusters (GCs) into gene cluster communities (GCCs) based on their shared distant homologies, which couldn't be detected with the sequence similarity approach. The GCC inference, described in the Methods-Cluster communities inference section, was implemented and tuned on the known sequence space, which is constrained by the domain architectures (DAs). Then, we used the information retrieved for the known sequence space to aggregate the unknown GCs. Since the number of DAs in the known GCs may be inflated due to the fragmented nature of metagenomic genes, a key step for the inference process was the retrieval of a set of non-redundant DAs (Methods - **A set of non-redundant domain architectures** section).

We reduced the complete set of 29,341 Pfam DAs found in the metagenomic dataset, to 23,681 non-redundant DAs, and the 38,765 Pfam DAs found in the genomic dataset to 38,060 non-redundant DAs.

To find how the different clusters aggregate at the DA level, we then applied a combination of HMM-HMM searches and community identification using the Markov Cluster Algorithm (MCL) (van Dongen & Abreu-Goodger, 2012) (see Methods - **Cluster communities inference)**. MCL is very sensitive to the inflation value, which determines the granularity of the partitioning. The results of our iterative approach are summarized in the radar plots of Supp. Figure 7-1. We determined the best inflation value at 2.2 for the metagenomic dataset, value corresponding to the radar plot with the largest area (Supp. Fig. 7-1A). This value is in agreement with the value empirically determined to be the optimal (van Dongen & Abreu-Goodger, 2012). The inference led to a set of 283,314 metagenomic GCCs out of ~2.9M GCs, with a reduction rate of 90% (Supp. Table 7-1A).

For the genomic dataset, we first identified the GCs with remote homologies to the metagenomic GCCs. To do this, we searched the genomic GC HMM profiles against the metagenomic ones, using HHblits (Remmert et al., 2012) (-n 2 -Z 10000000 -B 10000000 -e 1). We assigned the genomic GCs sharing a HHblits probability ≥ 50% and a bidirectional coverage > 60% to the respective metagenomic GCCs. We processed the remaining genomic GCs through the GCC inference workflow. We determined the best inflation value at 2.5 (Supp. Fig. 7-1B), which led to the inference of a total of 496,930 GCCs, with a reduction rate of 79% (Supp. Table 7-1B). The numbers of identified cluster GCCs for each category are shown in Supplementary Table 7-1.

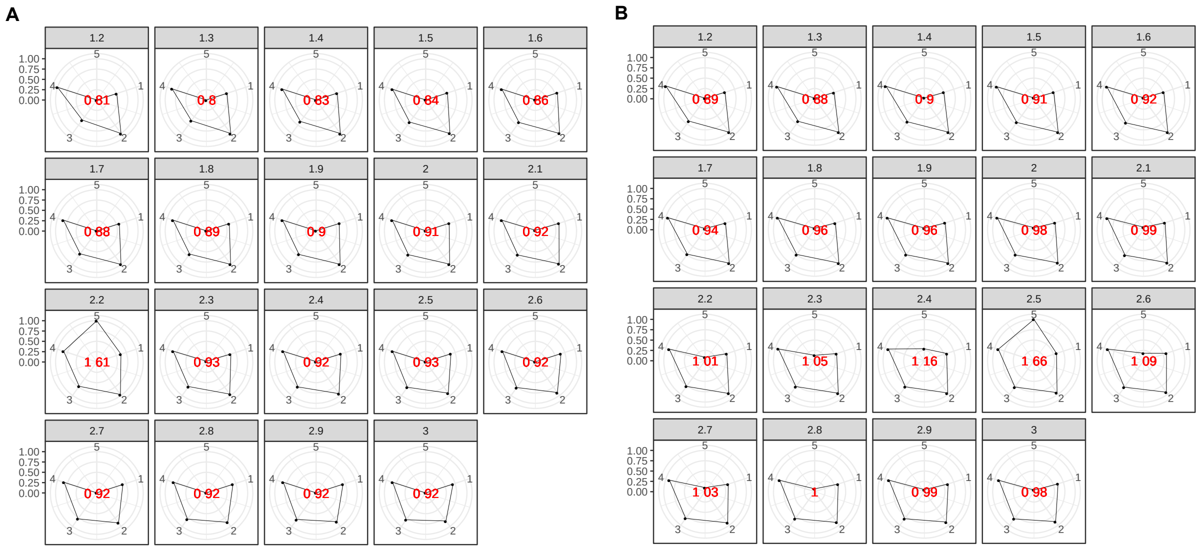


**Supplementary Figure 7-1.** Radar plots used to determine the best MCL inflation value for the partitioning of the K into cluster components. The plots were built using a combination of five variables: 1=proportion of clusters with one component and 2=proportion of clusters with more than one member, 3=clan entropy (proportion of clusters with entropy = 0), 4=intra HHblits-Score/Aligned-columns (normalized by the maximum value), and 5=number of clusters (related to the non-redundant set of DAs). (A) Metagenomic dataset. (B) Genomic dataset.

**Supplementary Table 7-1.** Number of gene clusters, cluster communities and reduction rate shown by functional category.

(A) Metagenomic dataset (MG)

| | K | KWP | GU | EU | Total |
|---|---|---|---|---|---|
| Clusters | 1,050,166 | 632,453 | 1,121,809 | 135,829 | 2,940,257 |
| Communities | 24,181 | 64,938 | 146,100 | 48,095 | 283,314 |
| Reduction (%) | 97.7 | 89.73 | 86.98 | 64.59 | 90.36 |

(B) Genomic dataset (GTDB)

| | K | KWP | GU | EU | Total |
|---|---|---|---|---|---|
| Clusters | 617,344 | 136,406 | 1,525,550 | 68,202 | 2,347,502 |
| Communities | 52,360 | 47,203 | 339,468 | 57,899 | 496,930 |
| Reduction (%) | 91.52 | 65.39 | 77.75 | 15.11 | 79.30 |

### Supplementary Note 8 - Gene cluster community validation

*The biological significance of the gene cluster communities (GCC) was tested by exploring their distribution within the phylogeny of proteorhodopsin and a set of ribosomal protein families.*

**Methods**

*Analysis of the GCC distribution within the proteorhodopsin phylogeny.*

We searched the proteorhodopsin (PR) HMM profiles from Olson et al.(Olson et al., 2018) against the K and KWP cluster consensus sequences, using the hmmsearch program of the HMMER software (version 3.1b2) (Finn et al., 2011). We filtered the results for alignment coverage > 0.4 and e-value ≥ 1e-5. The filtered results were placed in the MicRhoDE PR tree (Boeuf et al., 2015) using pplacer (Matsen et al., 2010). Then we placed the query PR sequences into the MicRhode (Boeuf et al., 2015) PR tree. We de-duplicated the placed queries with CD-HIT (v4.6) (Li & Godzik, 2006) and we cleaned them from sequences with less than 100 amino acids using SEQKIT (v0.10.1) (Shen et al. 2016). Next, we calculated the best substitution model using the EPA-NG modeltest-ng (v0.3.5) (Barbera et al., 2019) and we optimized the MicRhoDE PR tree initial parameters and branch lengths using RAxML (v8.2.12) (Stamatakis, 2014). Afterward, we incrementally aligned the query PR sequences against the PR tree reference alignment using the PaPaRA (v2.5) software (Berger & Stamatakis, 2012). We divided the query alignment and the reference alignment using EPA-NG –split v0.3.5. We combined the PR tree with the related contextual data and the tree alignment, into a phylogenetic reference package using Taxtastic (v0.8.9), and we placed the PR query sequences in the tree using pplacer (v1.1.alpha19-0-g807f6f3) (Matsen et al., 2010) with the option -p (–keep-at-most) set to 20. We grafted the PR tree with the query sequences using Guppy, a tool part of pplacer. 3. As the last step, we assigned the PR Supercluster affiliation to the query sequence, transferring the annotation of its closest relative in the MicRhoDE tree (Boeuf et al., 2015) the R packages APE v5.3 and phanghorn v2.5.3 (Schliep, 2011).

Furthermore, we aligned the query sequences annotated as viral to the six viral PRs from Needham et al. 2019 (Needham et al., 2019), using Parasail (Daily, 2016) (-a sg_stats_scan_sse2_128_16 -t 8 -c 1 -x). We then built a sequence similarity network (SSN) using the sequence similarity values to weight the graph edges.

*Analysis of standard and high-quality GCCs distribution within ribosomal protein families.*

As an additional evaluation, the distributions of standard GCCs and HQ GCCs within ribosomal protein families were investigated and compared. The ribosomal proteins used for the analysis were obtained combining the set of 16 ribosomal proteins from Méheust et al. (Méheust et al., 2019) and those contained in the collection of bacterial single-copy genes of anvi'o (Eren et al., 2015), that can be downloaded from (https://github.com/merenlab/anvio/blob/master/anvio/data/hmm/Bacteria_71/genes.txt).

#### Results

The results of both distribution analyses are shown in Figure 2D and 2C, respectively, and described in the main text.

We found 63 of the viral genes placed in the PR tree showing an average similarity of 50% with the viral PR of Needham et al. (Needham et al., 2019) (Supp. Table 8-1). Additionally, we found two genes (from two TARA samples: TARA_093_SRF_0.22-3 and TARA_145_SRF_0.22-3) sharing a similarity of 100% with one of the Needham et al. PRs (ChoanoV2_VirRyml_1). These genes, however, were not placed in the PR tree.

**Supplementary Table 8-1.** Sequence similarity values between viral genes and Needham et al. viral PRs. (Supplementary_tables_1.xlsx).

### Supplementary Note 9 - HMM-HMM homology network weighting metrics

*Validation of the edge weight metrics used for the gene cluster homology network community inference.*

#### Methods

A critical step in the gene cluster community (GCC) inference relies on the determination of the edge weights for the GC HMM-HMM network. We tested two possible metrics to weight the GC homology network resulting from the all-vs-all HMM GC comparison with HHblits(Remmert et al., 2012): (1) the ratio between the HHblits score and the number of aligned columns (*HHblits-Score/Aligned-columns*), metric chosen in this paper; (2) the maximum(HHblits-probability x coverage), weight used in Méheust et al. (2019) (Méheust et al., 2019). In addition, we tested the two different metrics using the ribosomal protein families as reference. For this second test, we filtered the GCCs for those annotated to the 16 ribosomal proteins used in Méheust et al. (Méheust et al., 2019), and those contained in the collection of bacterial single-copy genes of Anvi'o (Eren et al., 2015), which can be downloaded from https://github.com/merenlab/anvio/blob/master/anvio/data/hmm/Bacteria_71/genes.txt. To then compare the two metrics, we used the functions of the R package *aricode* (https://github.com/jchiquet/aricode)(Vinh et al., 2009), which allow comparisons between clustering methods.

#### Results

The results of the test of the different HHblits metrics used to weight the GC homology network are shown separately in Supp. Figure 9-1 and the comparison in Supp. Figure 9-2. Both metrics present a very different behavior (Supp. Fig. 9-1), the metric used in Méheust et al. is rescaling the *HHblits-probability* (Supp. Fig. 9-2). While the *HHblits-probability* is useful for deciding if two HMMs are reliable homologs, it is not suitable for measuring similarities due to its dependence on the length of the alignment. On top of this, we can see how the *HHblits-Score/Aligned-columns* values present a similar and more homogenous distribution in all four categories, being more suitable for the MCL clustering.

Overall, our approach generated fewer GCCs, as can be observed in Supp. Figure 9-3. Our clustering was found closer to the "*ground truth*" represented by the ribosomal protein families compared to the partitioning proposed by Méheust et al. The results from the comparison between the two clustering approaches and the ribosomal protein reference are reported in Supplementary Table 9-1.

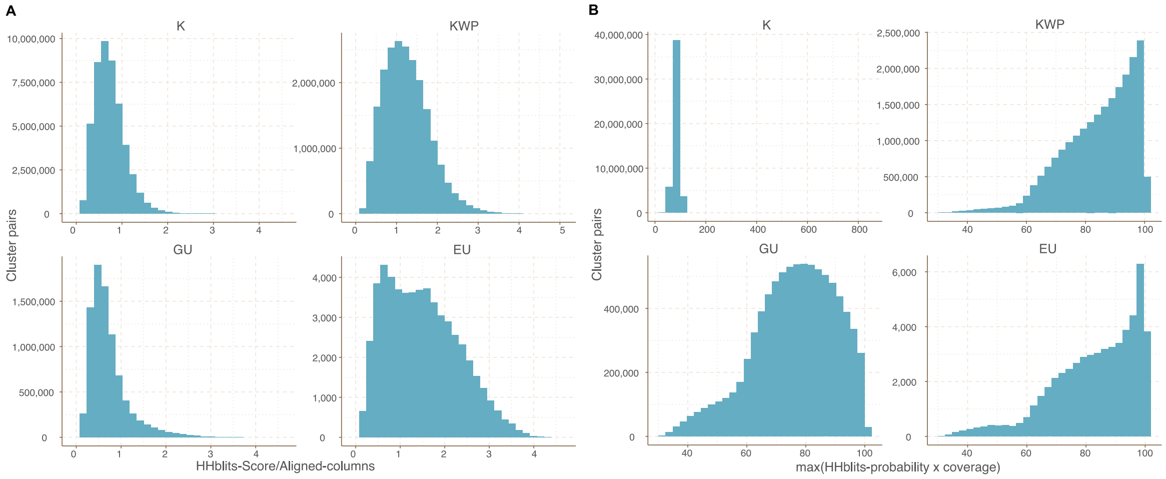


**Supplementary Figure 9-1.** Cluster pairs distribution based on the metrics used to weight the gene cluster HMM-HMM homology network. (A) HHblits-Score/Aligned-columns (Vanni et al.). (B) maximum(HHblits-probability x coverage) (Méheust et al.).

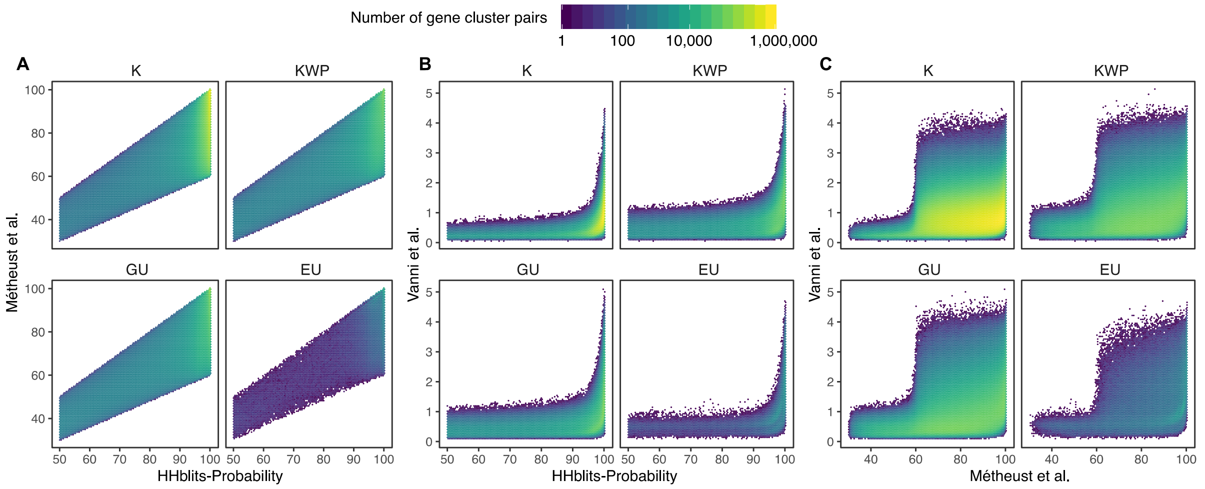


**Supplementary Figure 9-2.** Determination of the edge-weight metrics for the GC HMM-HMM homology network. We tested the metrics used in Méheust et al. and this paper (Vanni et al.). The correlations between metrics are shown per functional category. The metric used by Méheust et al. corresponds to the maximum(HHblits-probability x coverage). The metric applied in this manuscript is *HHblits-Score/Aligned-columns*. (A) Comparison between the metric of Méheust et al. and the HHblits-Probability. (B) Comparison between the metric used in this manuscript and the HHblits-Probability. (C) Comparison between the metric used in this manuscript and the metric of Méheust et al.

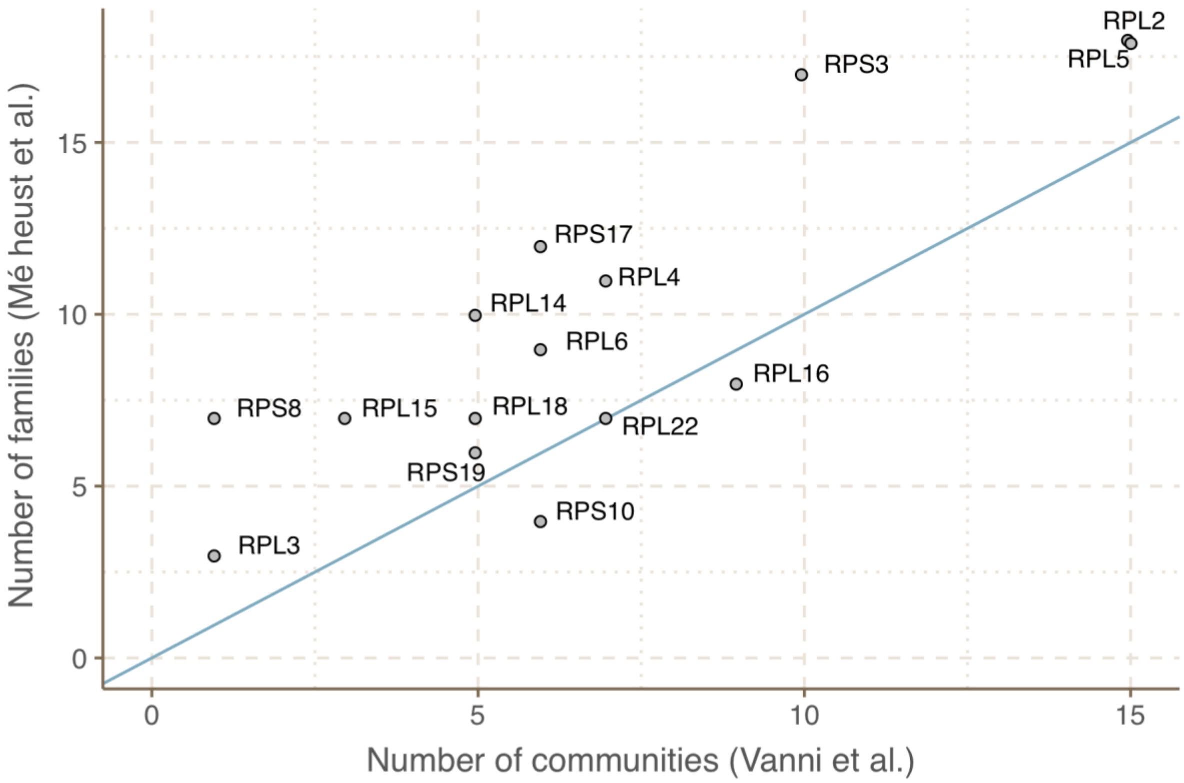


**Supplementary Figure 9-3.** Agreement between the number of communities within ribosomal protein families between our approach and the one described in Méheust et al.

**Supplementary Table 9-1.** Measures of similarity between the community inference approach proposed in this paper, the one used in Méheust et al. and the "ground truth" represented by the ribosomal protein families.

| | Vanni et al. vs Meheust et al. | Vanni et al. vs ribosomal families | Meheust et al. vs ribosomal families |
|---|---|---|---|
| ARI | 0.915 | 0.944 | 0.906 |
| AMI | 0.928 | 0.916 | 0.878 |
| NVI | 0.101 | 0.0858 | 0.124 |
| NID | 0.0717 | 0.0841 | 0.122 |
| NMI | 0.928 | 0.916 | 0.878 |

**Note:** ARI=Adjusted Rand Index; AMI=Adjusted Mutual Information; NVI=Normalized Variation Information; NID=Normalized Information Distance; NMI=Normalized Mutual Information.

#### Supplementary Note 10 - Singletons effect on the sequence space diversity

*Insights into the metagenomic and genomic singletons and their influence on the gene cluster rate of accumulation.*

Singletons represent a significant fraction in both the metagenomic (60%) and genomic (55%) datasets. Although we discarded them from the primary analyses presented in this paper, we analyzed their composition in terms of functional categories. The analysis steps are described for the metagenomic singletons in Supp. Note 1, and, after the integration, we applied the same steps to the genomic singletons (Supp. Table 10-1). As shown in Supp. Note 1, the metagenomic singletons are highly represented by EU genes, while in the genomes we observed the majority of the singletons shared between GU and EU. In general, the singletons are characterized by a high percentage of genes of unknown function.

We tested the singletons role in the rate of accumulation of GCs and GCCs as a function of the number of genomes and metagenomes, as shown in Figure 3C and 3D (to be compared with Supp. Fig. 5A and 5B). For the metagenomic collector curves, we included only the singletons with a sample abundance of 8.36. This value corresponds to the mode sample abundance of the set of metagenomic singletons that became clusters with more than ten genes after the integration of the genomic data.

We observed that, excluding the 19,911,324 singletons from the metagenomic dataset, the accumulation curves of the GCs flatten and approach a plateau. The same effect is observed, excluding the set of 5,558,438 singletons from the genomic dataset (Supp. Fig. 5B; Supp. Table 10-2).

**Supplementary Table 10-1.** Number of genomic singletons per functional category.

| | K | KWP | GU | EU |
|---|---|---|---|---|
| Genes | 473,460 | 896,127 | 2,528,370 | 1,660,481 |

**Supplementary Table 10-2.** Minimum slope values for the collector curves.

(A) Excluding singletons. In parenthesis, the number of genomes or metagenomes for the first occurrence of slope < 1

| Gene Clusters | Gene cluster Communities |
|---|---|

| | metaG | GTDB | metaG | GTDB |
|---|---|---|---|---|
| Known | 209.235 | 6.556 | 0.1344 (440) | 0.07 (15,120) |
| Unknown | 374.5147 | 5.851 | 0.1375 (600) | 0.621 (27,690) |

(B) Including singletons (with a mode abundance in the samples of 8.36).

| | Gene Clusters | |
|---|---|---|
| | metaG | GTDB |
| Known | 1329.489 | 66.063 |
| Unknown | 4843.570 | 158.891 |

### Supplementary Note 11 - Coverage of external databases

*Analysis of the coverage, by our metagenomic dataset, of seven external microbial gene and gene cluster datasets.*

**Methods**

We searched seven different state-of-the-art databases against our dataset of cluster HMM profiles. The different profile searches were all performed using the MMSeqs2 (version 8.fac81) *search* program (Steinegger & Söding, 2017), setting an e-value threshold of 1e-20 and a query coverage threshold of 60% (-e 1e-20 --cov-mode 2 -c 0.6). We kept the hits within 90% of the log10(best-e-value). Then we applied a majority vote function to retrieve the consensus functional category (K, KWP, GU or EU) for each search hit. In the end, the results were sorted by the lowest e-value and the largest query and target coverage to keep only the best hits.

We applied the described method to the following datasets: the Families of Unknown Functions (FUnkFams) (61,970 genes) (Wyman et al., 2018), the Pacific Ocean Virome (POV) (4,238,638 genes) (Hurwitz & Sullivan, 2013) and the Tara Ocean Virome (TOV) (6,642,187 genes) (Brum et al., 2015). The Genome Taxonomy Database (GTDB) (93,723,190 archaeal and bacterial genes) (Parks et al., 2018). The *MGnify* proteins from the EBI metagenomics database (release 2018_09)(Mitchell et al., 2018) (843,535,611 genes). The manually curated collection of 957 MAGs from TARA metagenomes (Delmont et al., 2017) (TARA MAGs) (2,288,202 genes), and the one made of 92 MAGs, from the fecal microbiota transplantation study (FMT MAGs) of Lee et al. (Lee et al., 2017) (188,983 genes). And also, the collection of unannotated genes with mutant phenotypes identified in Price et al. 2018 (Price et al., 2018) (37,684 mutant genes).

**Results**

We found our metagenomic GCs in all the main biomes defined by EBI metagenomics (Supp. Fig. 6), with an overall coverage of 74% of the MGnify peptides (Supp. Fig. 11-1). Our GCs also covered 62% of the FUnkFam genes of Wyman et al.; 70% of the GTDB genes; and 85% of the genes tested for mutant phenotypes in Price et al.. We also covered 50% of the Pacific Ocean Virome proteins, and 77% of the TARA Ocean Virome proteins, for overall coverage of 70% of the selected viral proteins. The majority of genes from both the FMT MAGs of Lee et al. and the TARA MAGs of Delmont et al., were found homologous to genes in our dataset (91% and 77% respectively). With the only exception of the FUnkFams, and the mutant genes, for which we did not find any homology to EU GCs, the other datasets reported homologies to clusters from all four functional categories. Moreover, we found that 20% of the Wyman et al

FUnkFams and 44% of the unknowns included in the RB-TnSeq experiments by Price et al., 2018 belong to the known sequence space (Supp. Table 11-1).

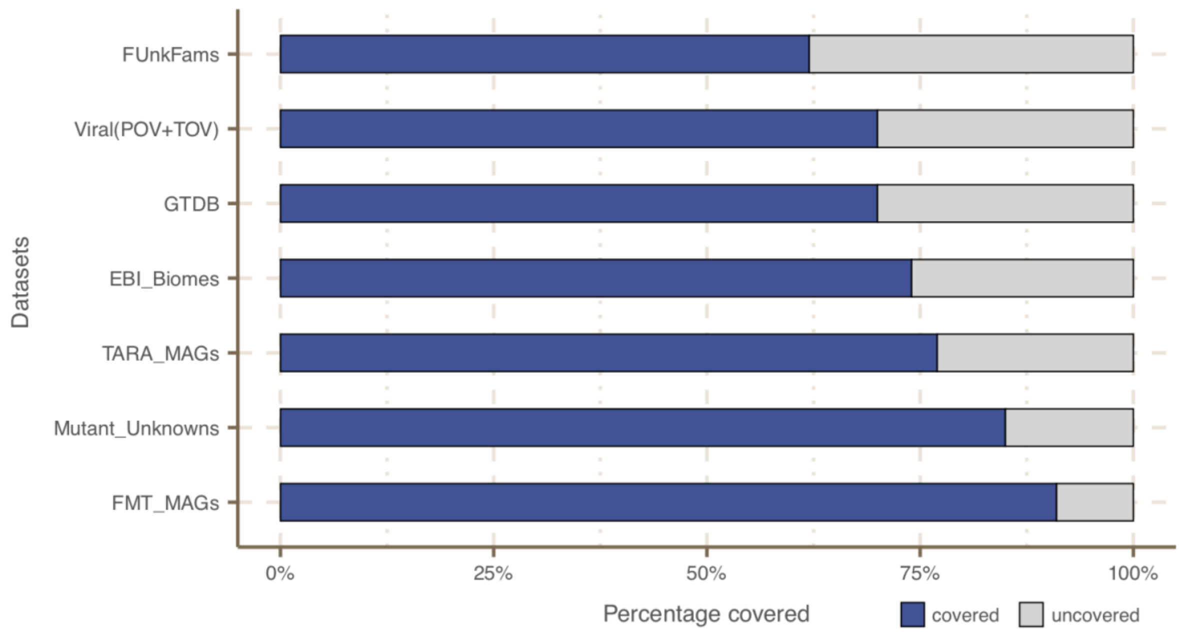


**Supplementary Figure 11-1.** Coverage of external datasets. The bar plot is showing the proportion of covered genes in each of the seven datasets that were screened against the metagenomic set of clusters' HMM profiles.

**Supplementary Table 11-1.** Re-classification of the unknowns identified in Wyman et al and Price et al.

| Study | Original unknown set | Covered fraction | Found as known | Found as unknown |
|---|---|---|---|---|
| Wyman et al. | 61,970 | 38,174 | 12,366 | 25,808 |
| Price et al. | 49,736 | 33,016 | 21,967 | 11,049 |

### Supplementary Note 12 - EU gene cluster in metagenome-assembled genomes

*Metagenome-assembled genomes (MAGs) as a resource to contextualize the environmental unknown gene clusters and cluster communities.*

Overall, the MG+GTDB integrated cluster dataset contains 204,031 EU gene clusters (GCs) (grouped in 103,195 cluster communities (GCCs)). The EUs are divided into 127,032 metagenomic, 70,470 genomic, and 9,024, both metagenomic and genomic GCs. The last two subsets contain 52,231 (26%) EU found in GTDB metagenome-assembled genomes (MAGs). To test whether we could also place the subset of metagenomic EU in the context of MAGs, we searched the GCs of this set against the manually curated TARA Ocean MAG collection from Delmont et al. (Delmont et al., 2017).
In addition, we deepened the investigation of the metagenomic EU subset, focusing on the GCCs found broadly distributed in metagenomes according to the results of Levin's niche breadth analysis (Fig. 4). The details of the metagenomic EU analysis are described below.

**Methods**

We searched the metagenomic EU GCs HMM profiles, obtained from the cluster MSA using the *hhmake* program of the HH-SUITE (Steinegger et al., 2019), against the set of 957 high-quality MAGs binned from the TARA Ocean prokaryotic dataset(Delmont et al., 2017). We performed the sequence-profile search using the MMSeqs2 *search* program (Steinegger & Söding, 2017), using -e 1e-20 --cov-mode 2 -c 0.6. We filtered the results to keep the hits within 90% of the log10(best-e-value). We applied a majority vote function to retrieve the consensus category for each hit. Then, we sorted the results by the smallest e-value and the largest query and target coverage to keep only the best hits. We then filtered the search results focusing on the broadly distributed EU GCs and GCCs. We retrieved MAG contigs containing the EU GCs and GCCs from the Anvi'o MAG profiles using the program *anvi-export-gene-calls* from Anvi'o v4 (Eren et al., 2015). We functionally annotated the contigs searching their genes against the Pfam database (v. 31.0) (Finn et al., 2016), using the *hmmsearch* program from the *HMMER* package (version: 3.1b2) (Finn et al., 2011), and complementing the search using Prokka (Seemann, 2014) in metagenomic mode. We then selected the contig with the lowest percentage of hypothetical proteins, and we extracted a region of 1kb surrounding the genes mapping to the EU GCCs.

**Results**

We found a total of 5,420 EU clusters homologous to 7,661 genes in the 691 TARA MAGs. These EU clusters belong to 4,365 GCCs. We kept only the 71 EU GCCs that showed a broad distribution in TARA samples. These GCCs contained 3,119 clusters and were found in 83 different TARA MAGs. Next, we examined the genomic neighborhood of the broad distributed EU on the MAG contigs. Investigating the genomic neighborhood can lead to the inference of a possible function of the EU. We selected the MAG most enriched with broadly distributed EU, which resulted in being the Atlantic North-West MAG "TARA_ANW_MAG_00076" (Supp. Fig. 12-1A). This MAG contains 23 EU (0.3%) of its genes. It belongs to the bacterial order of *Flavobacteriales*. Of its 1,283 contigs, 317 include at least one EU. We functionally annotated these contigs with Prokka (and Pfam). Then, we sorted the contigs based on the proportion of genes annotated to hypothetical or characterized proteins, as shown in Supp. Figure 12-1B. The presence of genes of known function around the EU contributes to prove that these unknown genes are part of a real contig, and possibly an operon. Therefore, we selected for exploration, the contigs with the highest proportion of characterized genes, "TARA_ANW_MAG_00076_000000000672", with 7 characterized genes out of a total of 13 annotated genes. The contig with the second least number of hypothetical proteins was "TARA_ANW_MAG_00076_000000001247", which contained nine characterized genes out of 20. The contig "TARA_ANW_MAG_00076_000000000672" is shown in Supp. Figure 12-1C and highlighted in red are the two predicted genes with significant homology to the EU GCs, members of the broadly distributed EU GCCs eu_com_769 and eu_com_5081. Within their genomic neighborhood, we observe genes relating to nucleotide metabolism, DNA repair and phosphate regulation/sensing, including dUTPase, phoH and protein RecA. Gene placement in prokaryotic genomes is not random. Genes are grouped to increase transcriptional efficiency to respond to stimuli in the environment. Therefore, we can hypothesize that these EU have functions related to their neighboring genes.

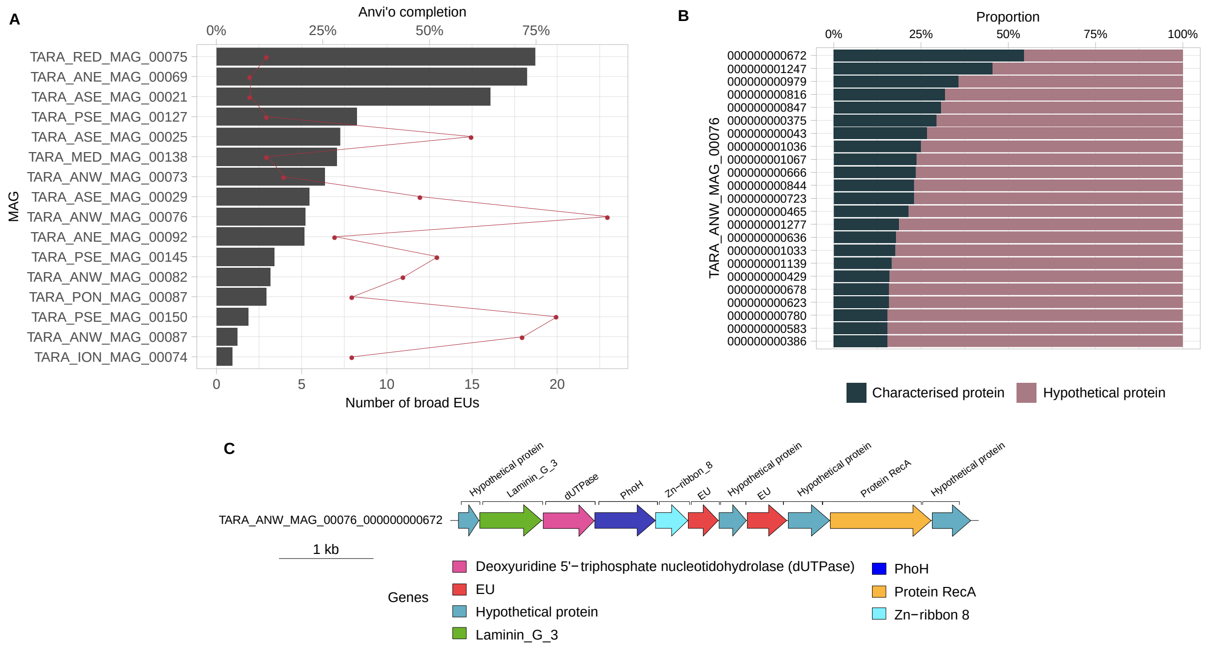


**Supplementary Figure 12-1.** (A) EU mapping on TARA MAGs results. Histogram of TARA MAG percent completeness (checkM). The red line represents the number of EU found in the MAGs. (B) Contigs from TARA MAGs TARA_ANW_MAG_00076 in descending order of highest proportion of non-hypothetical gene content. (C) EU communities in the context of a MAG contig. Contig genomic neighborhood around two potential EU communities.

### Supplementary Note 13 - Archaea gene cluster phylogenomic analysis

*Gene clusters phylogenetic analysis - results for the archaeal genomes.*

In the main text are shown the results for the gene clusters (GCs) phylogenetic analyses (clusters phylogenetic conservation and specificity) for the GTDB bacterial genomes. The same methods/analyses were applied for the archaeal genomes, and the results are presented here.

Out of the 230,340 GCs found in GTDB archaeal genomes, we identified 48,518 lineage-specific GCs (precision and sensitivity both ≥95% (Mendler et al., 2019)). As seen for the Bacteria in Figure 5A, the number of known and unknown archaea lineage-specific GCs increases with the Relative Evolutionary Distance (Parks et al., 2018), with the differences between the known and the unknown fraction starting to be evident at the Family level (Supp. Fig. 13-1A). The number of unknown lineage-specific GCs for Family, Genus and Species are 2,937, 12,966 and 21,002 respectively (Supp. Tale 13-1). A total of 34,893 GCs were phylogenetically conserved ($P < 0.05$), where 19,693 were known GCs and 15,200 were unknown GCs. Overall, the unknown GCs are more phylogenetically conserved than the known GCs (Supp. Fig. 13-1B, $p < 0.0001$). However, considering only the lineage-specific clusters, we observe the opposite, the unknown GCs result in less phylogenetically conserved (Supp. Fig. 13-1B). The GTDB archaeal genomes were also screened for prophages. In total, we identified 2,082 lineage-specific GCs in prophage genomic regions, and 86% of them resulted in clusters of unknown function (Supp. Fig. 13-1C). To identify archaeal phyla enriched in unknown GCs, we partitioned the phyla based on the ratio of known to unknown GCs and vice versa (Supp. Fig. 13-1D). We observed the same pattern found for bacterial phyla in Figure 5D, where the archaeal phyla with a larger number of MAGs are enriched in GCs of unknown function (Supp. Fig. 13-1D).

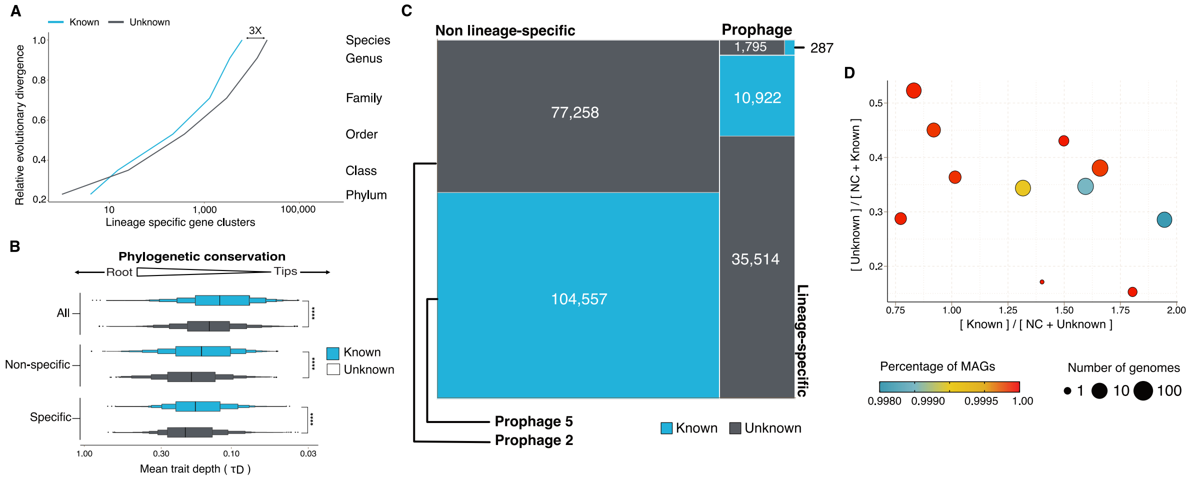


**Supplementary Figure 13-1.** Phylogenomic exploration of the unknown sequence space in Archaea. (A) Distribution of the lineage-specific gene clusters by taxonomic level. Lineage-specific unknown gene clusters are more abundant at the lower taxonomic levels (genus, species). (B) Phylogenetic conservation of the known and unknown sequence space in 1,569 archaeal genomes from GTDB. We calculated the mean trait depth ($\tau_D$) with the consenTRAIT algorithm and the lineage specificity using the F1-score approach from (Mendler et al., 2019). We observe differences in the conservation between the known and the unknown sequence space for lineage- and non-lineage-specific gene clusters (paired Wilcoxon rank-sum test; all p-values < 0.0001). (C) The majority of the lineage-specific clusters are part of the unknown sequence space, being a small proportion found in prophages present in the GTDB genomes. (D) Known and unknown sequence space of the 1,569 GTDB archaeal genomes grouped by archaeal phyla. Phyla are partitioned based on the ratio of known to unknown gene clusters and vice versa from the set of genomes. Phyla enriched in Metagenomic assembled genomes (MAGs) have a higher proportion in gene clusters of unknown function.

**Supplementary Table 13-1.** Number of phylogenetic conserved and lineage-specific GCs in the GTDB archaeal phylogeny. (Supplementary_tables_1.xlsx).

### Supplementary Note 14 - *Cand.* Patescibacteria lineage-specific gene clusters analysis

*The investigation of the lineage-specific clusters was deepened, focusing on those specific to the Cand. Patescibacteria phylum (former Candidate Phyla Radiation-CPR) and analyzing their cluster distribution in both the Human and marine (TARA and Malaspina) metagenomes.*

We found two GU clusters phylum-specific, and a total of 54,343 clusters of unknown function, lineage-specific within the *Cand.* Patescibacteria phylum (Supp. Table 14-1). The majority of this phylum members are particularly poorly understood microorganisms, mostly due to undersampling and the incompleteness of the available genomes. Therefore, we decided to investigate the distribution in the human and marine (TARA and Malaspina) metagenomes of all the clusters lineage-specific inside the *Cand.* Patescibacteria phylum (Supp. Fig. 14-1A).
We chose to have a closer look at the class of *Gracilibacteria,* which shows to be present in both human and marine environments. The first genome for this class was retrieved in a hydrothermal vent environment in the deep sea (Rinke et al., 2013). The same organisms were then also identified in an oil-degrading community (Rinke et al., 2013; Sieber et al., 2019) and as a part of the oral microbiome (Espinoza et al., 2018). As shown in Supp. Figure 14-1B, we found both known and unknown clusters lineage-specific to this class, distributed in human and marine metagenomes. Among these clusters, we observed cases of environment specificity. For instance, three clusters of unknowns were found exclusive to HMP samples. These clusters could be proposed as novel targets for human-health study since *Gracilibacteria* was found enriched in healthy individuals (Espinoza et al., 2018). We also observed lineage-specific clusters of known and unknown functions specific to the marine environment.

**Supplementary Table 14-1.** Number of lineage-specific clusters within the *Cand.* Patescibacteria phylum, at different taxonomic levels, subdivided by cluster categories.

| Taxonomic level | K | KWP | GU | EU |
|---|---|---|---|---|
| Phylum | 1 | 0 | 2 | 0 |
| Class | 11 | 0 | 6 | 0 |
| Order | 41 | 1 | 104 | 0 |
| Family | 452 | 9 | 1,443 | 13 |
| Genus | 625 | 98 | 6,649 | 338 |
| Species | 4,116 | 818 | 42,710 | 3,078 |

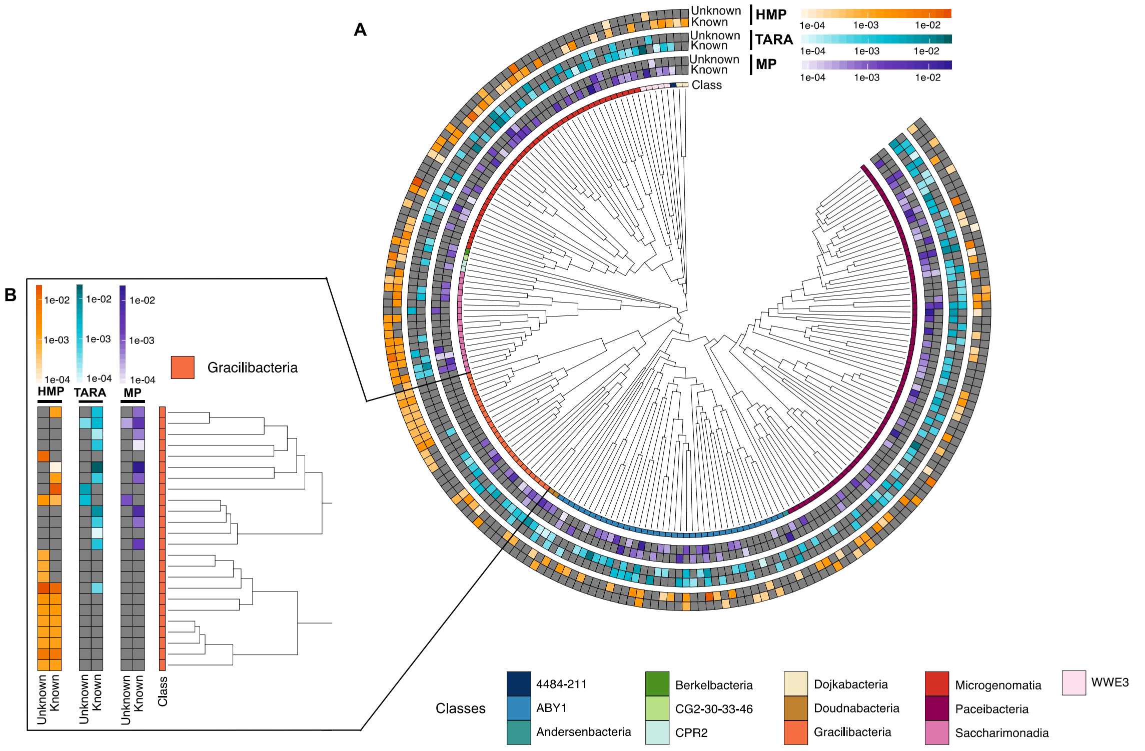


**Supplementary figure 14-1.** *Cand.* Patescibacteria metagenomic lineage-specific clusters. (A) Phylogenetic tree of *Cand.* Patescibacteria genera, colored by classes. The heatmaps around the tree show the proportion of lineage-specific gene clusters of knowns and unknowns in the metagenomes from TARA, Malaspina and the HMP. (B) Metagenomic lineage-specific clusters in the class of *Gracilibacteria*.
